# The Ca^2+^ Channel CYCLIC NUCLEOTIDE GATED CHANNEL13 (CNGC13) regulates systemic wound signalling and immunity against Spodoptera herbivory

**DOI:** 10.1101/2025.06.16.660006

**Authors:** Ramgopal Prajapati, Misha Kumari, Yashaswi Singh, Mahendra Pawar, Bianca Maria Orlando Marchesano, Jeet Kalia, Alex Costa, Jyothilakshmi Vadassery

**Affiliations:** BRIC-National Institute of Plant Genome Research (NIPGR), Aruna Asaf Ali Marg, New Delhi 110067, India; Department of Biological Sciences, Indian Institute of Science Education and Research (IISER) Bhopal, Madhya Pradesh, India; Department of Biosciences, University of Milan, via Celoria 26, 20133 Milano, Italy; Institute of Biophysics, National Research Council of Italy (CNR), Milan, 20133, Italy

**Keywords:** Herbivory, Plant Immunity, Calcium Channels, CNGC, Jasmonates, Glucosinolate This PDF file includes:

## Abstract

Insect herbivory elicits rapid and complex signalling responses in plants, including membrane depolarization, cytosolic Ca^2+^ elevation and activation of jasmonate-mediated defense pathways in local and systemic leaves. These early events are orchestrated by calcium-permeable channels and regulatory components that generate stimulus-specific cytosolic Ca^2+^ signatures. However, the molecular mechanisms that generate long-distance Ca^2+^ propagation and systemic immune activation on herbivory remain poorly defined. Here, we identify CYCLIC NUCLEOTIDE GATED CHANNEL13 (CNGC13) as a novel plasma membrane-localized, phloem-expressed Ca^2+^ channel that is essential for systemic wound signalling and defense against lepidopteran pest, *Spodoptera litura* in *Arabidopsis thaliana*. CNGC13 is rapidly induced by wounding and herbivory, mediates cytosolic Ca^2+^ influx, and is required for efficient propagation of Ca^2+^ signals from wounded to distal leaves. Loss of CNGC13 compromises the thioglucosidase-dependent breakdown of aliphatic glucosinolates into aglucones/isothiocyanates within the vascular tissue, which are essential for wounding-induced systemic Ca^2+^ wave propagation. Consequently, *cngc13* mutants display impaired systemic jasmonate accumulation in distal leaves, reduced glucosinolate levels and enhanced herbivore susceptibility. CNGC13 is also required for *At*Pep-induced Ca^2+^ elevation and Reactive Oxygen Species (ROS) signalling and interacts with the receptor kinase, PEPR2, implicating it in DAMP-mediated immune responses. Although CNGC13 and CNGC19 function cooperatively in herbivore defense, they do not physically interact, suggesting independent but converging roles. Our study identifies CNGC13 as a vasculature-localized Ca^2+^-permeable channel that functions downstream of PEPR2 receptor, to sustain systemic signalling, operating in a distinct yet converging pathway alongside known glutamate receptor-like channels during long-distance wound responses.

**Significance:** Plants must detect and respond to insect herbivory rapidly by activating systemic immunity to trigger defense responses. Calcium (Ca^2+^) signalling plays a central role in this defense network, yet the identities and specific functions of Ca^2+^ channels involved in long-distance herbivory signalling remain poorly understood. Here, we identify the plasma membrane-localized channel CYCLIC NUCLEOTIDE GATED CHANNEL13 (CNGC13) as a key integrator of mechanical damage perception, ricca factor-mediated systemic signalling, and jasmonate-dependent defense against *Spodoptera* herbivory in *Arabidopsis*. CNGC13 facilitates systemic Ca^2+^ wave propagation, activates jasmonate and glucosinolate biosynthesis, and promotes the perception of damage-associated molecular pattern (DAMP) molecules to confer herbivore resistance. This work uncovers a previously uncharacterized calcium channel that plays a crucial dual role in systemic wound signalling.

## Introduction

Plant–insect interactions, among the most widespread biotic interactions in nature, are responsible for approximately 20% of global annual crop losses and are projected to intensify further with rising global temperatures (Lehmann et al., 2020). In response to chewing insect attack, plants recognize mechanical damage and specific chemical cues in herbivore oral secretions, damage-associated molecular patterns (DAMPs) and herbivore-associated molecular patterns (HAMPs) respectively, to activate defense signalling cascades (Snoeck et al., 2022). Among DAMPs, peptides (*At*Peps) and extracellular ATP (*e*ATP) are well-characterized signals that trigger cytosolic Ca^2+^ elevations and defense-related phytohormone production on herbivory (Kundu et al., 2025; Klauser, 2015). Rupture of veins due to herbivory results in a loss of xylem tension and phloem pressure which allows uptake of wound-derived DAMPs or herbivore-associated chemical signals into damaged vessels, facilitating their long-distance transport throughout the plant (Farmer et al., 2020; Gao et al., 2023). Perception of these HAMPs and DAMPs is mediated by pattern recognition receptors (PRRs) localized at the plasma membrane. These molecular patterns and their cognate receptor complexes form a damage perception system that generates specific immune signals (Koster *et al*., 2022).

Wounding rapidly initiates a cascade of biochemical and biophysical events, including changes in turgor pressure, slow wave potential and cytosolic Ca^2+^ levels which propagate from the local wound site to distant, unwounded tissues (Farmer et al., 2020; Hedrich et al., 2023). Among these, transient elevations in cytosolic Ca^2+^ concentration are among the earliest and most crucial signals, propagating across tissues and triggering local and systemic defense responses. Upon perception of different stimuli, the cytosolic influx of Ca^2+^ is mediated by several classes of ion channels, including cyclic nucleotide-gated channels (CNGCs), glutamate receptor-like channels (GLRs), two-pore channels (TPCs), mechanosensitive channels (MCAs), and reduced hyperosmolality-induced [Ca^2+^]_i_ increase (OSCAs) (Yuan et al., 2014; Demidchik et al., 2018). In particular, in response to wounding, these Ca^2+^ signals are coupled with leaf surface potential depolarization, hydraulic pressure changes, and phytohormone biosynthesis in vasculature to facilitate a coordinated systemic response (Nguyen et al., 2018; Grenzi et al., 2023). One of the primary downstream consequences of Ca^2+^ signalling is the activation of jasmonic acid (JA) biosynthesis, a key phytohormone involved in defense against chewing pests like *Spodoptera litura*. The events leading to jasmonate production in and near the wound site are highly complex and intertwined with electrical and Ca^2+^ signal propagation (Mousavi et al., 2013; Farmer et al., 2014; Ngyuen et al., 2018). Secondary metabolites, which act downstream of phytohormones, also play essential roles in plant defense. Glucosinolates (GS) are well-characterized secondary metabolites in *Arabidopsis* that remain biologically inert until tissue damage brings them into contact with the myrosinase enzymes/ THIOGLUCOSIDE GLUCOHYDROLASE 1 (TGGs). This contact firstly triggers their hydrolysis leading to the formation of an unstable intermediate and finally to a variety of toxic breakdown products like isothiocynates, that act as a feeding deterrent against generalist herbivores (Textor and Gershenzon, 2009). A recent study places localized glucosinolate-myrosinase system, much early in immune signalling with a role in leaf-to-leaf wound-response signalling through mobile TGG proteins (ricca factors). Mobile TGGs are, at least in part, responsible for inducing the slow wave potential and Ca^2+^ wave propagation via aglucone mediated membrane depolarization in Arabidopsis (Gao et al., 2023).

The systemic wound response on herbivory relies on both chemical and electrical signals that travel through the vasculature, where jasmonate biosynthesis occurs primarily in the xylem parenchyma cells (Chauvin et al., 2013; Farmer et al., 2014). This complex signalling involves a coordinated network of herbivory-induced cues and multiple classes of ion channels that operate across cellular compartments and tissue layers. Local and systemic increases in apoplastic glutamate occur in response to wounding and function as (DAMPs), that activate glutamate receptor-like channels (GLRs), that mediate extracellular glutamate-induced systemic Ca^2+^ waves (Toyota et al., 2018; Alfieri et al., 2020; Grenzi et al., 2023). Notably, *glr3.3 glr3.6* double mutants exhibit impaired propagation of both electrical and cytosolic Ca^2+^ signals between leaves, and subsequent lack of jasmonate accumulation in systemic leaves, underscoring their essential role in wound-induced long-distance signalling (Mousavi et al., 2013, Ngyuen et al., 2018; Toyota et al., 2018). GLR3.3 localises to phloem sieve elements, whereas GLR3.6 to xylem contact cells (Nguyen et al., 2018). The vacuolar channel TPC1 also contributes to systemic Ca^2+^ elevations on wounding and aphid feeding (Kiep et al., 2015; Vincent et al., 2017). The phloem-expressed Ca^2+^ channel CNGC19 mediates rapid wound-induced Ca^2+^ influx in local leaves and acts as a positive regulator of immunity against *Spodoptera litura* by modulating aliphatic glucosinolate biosynthesis (Meena et al., 2019). The mechanosensitive anion channel MSL10, expressed in both xylem and phloem, responds to membrane tension and contributes to wound-induced electrical and Ca^2+^ signalling (Moe-Lange et al., 2021). Apart from conventional ion channels, other Ca^2+^ signalling components also shape the pathway. These include the P-type H[-ATPase AHA1, which is required for the repolarization phase of the wound-induced slow wave potential and thereby modulates signal duration and downstream systemic jasmonate production (Kumari et al., 2019). Annexin (ANN1) plays an important role in activation of systemic defense in plants on herbivory by *Spodoptera littoralis* (Malarba et al., 2021). Systemic Ca^2+^ waves are known to rely not only on active ion channel-mediated propagation, but also on the diffusion and bulk flow of amino acids (glutamate) through the vasculature (Bellandi et al., 2022). Together, these findings suggest that herbivory-induced Ca^2+^ dynamics in local and systemic leaves are orchestrated by a suite of ion channels and signalling components acting across cellular compartments and tissue layers.

The interplay between Ca^2+^ signalling, jasmonic acid accumulation, and the production of defense metabolites highlights the complexity of the plant’s defense network. While the role of GLRs in systemic Ca^2+^ signal propagation is well established, the involvement of other Ca^2+^ channels and DAMPs, particularly in the context of herbivory, remains poorly understood. How plants integrate vascular damage-derived cues with Ca^2+^ channel activity and receptor-mediated signalling to coordinate long-distance Ca^2+^ waves and systemic defense responses are still largely unknown. Plant cyclic nucleotide-gated channels (CNGCs) are plasma membrane-localized, non-selective cation channels crucial for Ca²⁺ signalling, but unlike their animal counterparts, they are not gated by cyclic nucleotides and instead produce inward-rectifying currents in response to hyperpolarizing voltages (Wang et al., 2025). The Arabidopsis CNGC family has 20 members subdivided into five groups (Mäser et al., 2001). *CNGC13* belongs to the group I, along with *CNGC10, CNGC11, CNGC12, CNGC3* and *CNGC1*. *CNGC11* and *CNGC12* have been intensively characterized to have role in plant defense against pathogen attack (Yoshioka *et al*., 2006; Moeder *et al*., 2011). The only known function of CNGC13 is in heavy metal ion transport in roots and mutations in *At*CNGC13 results in reduced lead accumulation (Moon et al., 2019). There are no reports on role of CNGC13 in plant immunity. This study identifies how Ca^2+^-permeable channel CNGC13 contributes to systemic wound signalling and defense activation in Arabidopsis, and how it integrates with known DAMP signalling pathways to coordinate long-distance immune responses against insect herbivory.

## Results

### *CNGC13* is activated in local and systemic leaf upon wounding and is a positive regulator of herbivory

To identify early-response genes involved in defense against *Spodoptera litura* herbivory in *Arabidopsis thaliana*, we compared Affymetrix microarray data from wounding combined with *S. litura* oral secretion treatment (W+OS) versus wounding with water (W+W) for 30 minutes (Meena et al., 2019). This analysis revealed significant upregulation of several cyclic nucleotide-gated channel (CNGC) genes, including CNGC19 (Meena *et al*., 2019) and a previously uncharacterized gene, CNGC13 (At4g01010). To validate these findings, we performed a simulated herbivory experiment and monitored *CNGC13* expression over time. We found that both W+W and W+OS treatments rapidly induced *CNGC13* expression within 15 minutes. While W+W triggered only a transient mild response, W+OS caused a sustained upregulation, with expression peaking at 60 minutes and remaining elevated up to 90 minutes (**Figure 1A**). We further assessed *CNGC13* expression following actual *S. litura* feeding. Expression increased approximately 5-fold at 30 minutes, peaked at 1 hour, and returned to basal levels by 24 hours (**Figure 1B**). To assess systemic induction, we wounded the petiole of leaf 8 and measured *CNGC13* expression in a distal, unwounded leaf (leaf 13). *CNGC13* was rapidly induced systemically, with expression detectable within 30 minutes, and declining by 90 minutes (**Figure 1C**). To find out the functional role of *CNGC13,* we performed an insect-no-choice assay by measuring the weight gain of *S. litura* on WT, *cngc13-1* (Salk_057742), *cngc13-2* (Salk_082668), *cngc13-3* (CRISPR line) and *coi1-16* (jasmonate receptor, positive control) mutant lines (**Supplementary Figure 1A, B**) upon 8 days of feeding. There was no visible growth phenotype difference observed between WT and *cngc13* mutant lines (**Figure 1D**). *S. litura* larvae fed on a larger leaf area and gained significantly more weight on *cngc13* mutant lines, as compared to WT (**Figure 1D, 1E).** Data suggests that *CNGC13* acts as positive regulator of plant defense against *S. litura* herbivory and the increase in larval weight is indicative of decreased defense in *cngc13* plants.

**Figure 1:**
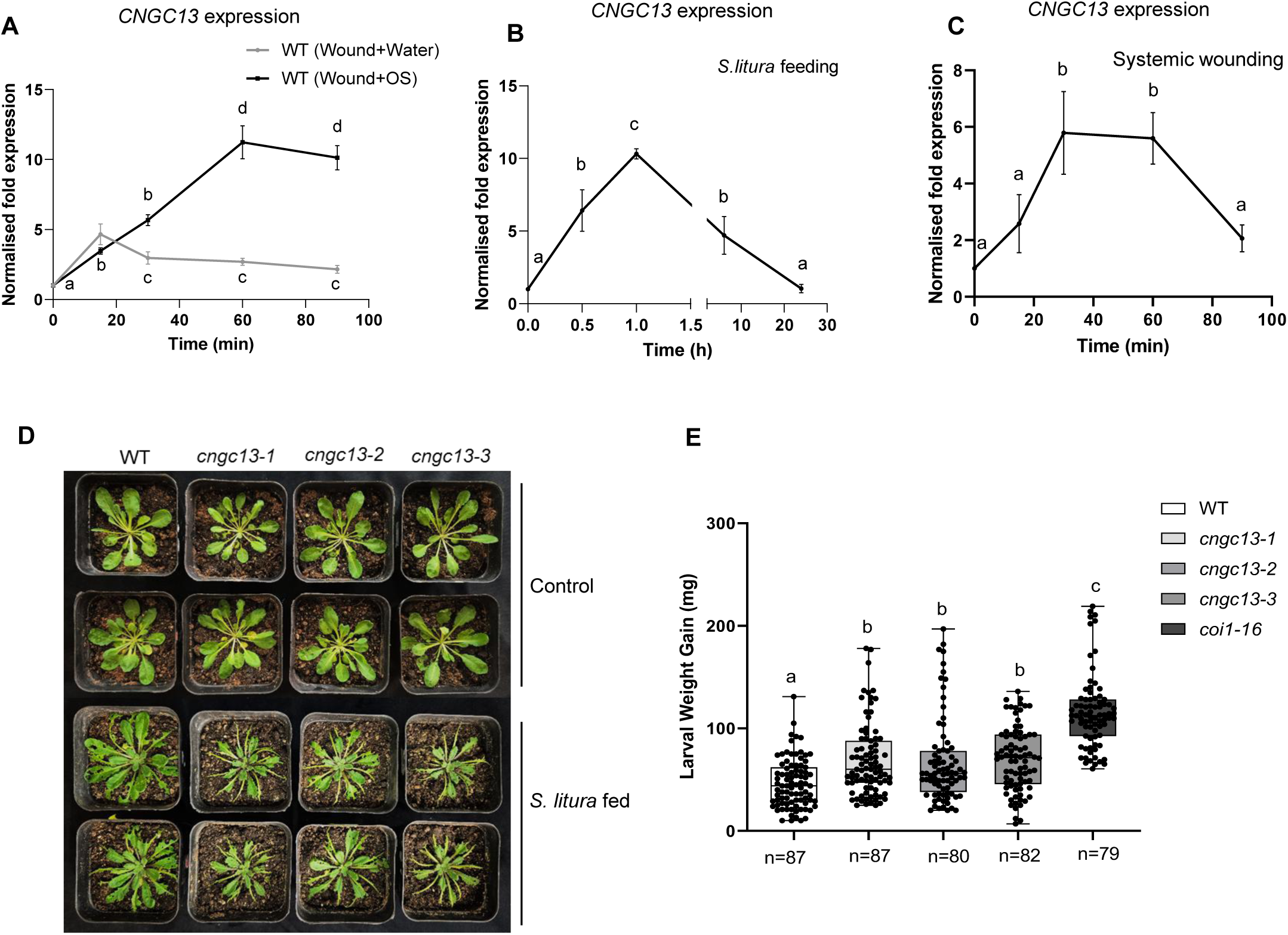
CNGC13 transcript is activated in local and systemic leaf upon wounding and herbivory and is a positive regulator of *Spodoptera litura* herbivory. (**A**) *CNGC13* transcript levels in *Arabidopsis* leaves upon W+W (grey) and W+OS (black) treatment for 15, 30, 60 and 90 min. The graphs show fold change (Mean ± SE) of mRNA level relative to untreated control. The figure shows data from the means of three independent experiments, n=4. Different letters indicate statistically significant differences among treatments, calculated by one-way ANOVA. (**B**) *CNGC13* transcript levels in *Arabidopsis* leaves after 0.5, 1, 6 and 24 h of *S. litura* feeding. The graphs show fold change (Mean ± SE) of mRNA level relative to untreated control. The figure shows data from the means of three independent experiments, n=4. Different letters indicate statistically significant differences among treatments, calculated by one-way ANOVA. (**C**) *CNGC13* transcript induction in the systemic leaf 13 at 0, 15, 30, 60 and 90 min after cutting the petiole of leaf 8. The graphs show fold change (Mean ± SE) of mRNA level relative to untreated control. The figure shows data from the means of three independent experiments with n = 6. Different letters indicate statistically significant differences among treatments, calculated by one-way ANOVA. (**D**) Phenotype of leaf area eaten upon feeding of *S. litura* larvae on Col-0, *cngc13-1* (Salk_057742), *cngc13-2* (Salk_082668) and *cngc13-3* (CRISPR line) after 8 days of feeding by second-instar *Spodoptera litura* larva in a no-choice-assay (**E**) Larval weight after no-choice assay on Col-0, *cngc13-1* (Salk_057742), *cngc13-2* (Salk_082668), *cngc13-3* (CRISPR line) and *coi1-16* (jasmonate receptor, positive control). *S. litura* 2^nd^ instar larvae were pre-weighed and three larvae were placed on per plant of each genotype. The total number of larvae weighed (n) is indicated in the bars. Box-whisker plots illustrating the larval weight measured after 8 d of feeding. Statistically significant differences between larvae fed on different genotypes were analyzed by one-way ANOVA with a with a post-hoc SNK test (*P* ≤ 0.05). Different letters indicate significant differences among treatments.

### *CNGC13* is expressed in phloem and plasma membrane localized

To investigate the spatial regulation of *CNGC13* in response to wounding, we analyzed its expression pattern in the leaf. CNGC13 promoter activity was barely detected in unwounded plants, but its activity was enhanced on wounding and was detected in primary, secondary and tertiary veins of wounded local and systemic leaf (**Figure 2A**). To verify tissue and cellular localization, we transformed the CNGC13 genomic DNA with a GFP tag under its native promoter (*pCNGC13::gCNGC13-GFP*) into WT and *cngc13-1* mutant. To identify the cell types involved in CNGC13-mediated signalling, we generated and analyzed Arabidopsis lines expressing *pCNGC13:gCNGC13-GFP* fusion constructs. Confocal imaging of primary root revealed GFP fluorescence in the vascular tissues (**Figure 2B**). To identify the exact localization in vasculature, aniline blue which stains callose deposition in phloem sieve plate was used. We find that CNGC13 accumulates in phloem as shown from the merged images (**Figure 2C**). We also confirmed this using *pCNGC13::GUS* lines and a longitudinal section of the petiole shows the GUS expression in phloem (**Figure 2D**). To further study the sub-cellular localization of CNGC13, we performed a co-localization experiment using *Nicotiana benthamiana* plants transiently co-transformed with 35S:CNGC13-YFP and plasma membrane marker, PM-mCherry. In *N. benthamiana* CNGC13 localized on the plasma membrane and overlays with PM-mCherry marker (**Figure 2E**). We also studied localization in the stable Arabidopsis lines expressing *pCNGC13::gCNGC13-GFP* fusion constructs. Using both root and leaf epidermal cells in these lines we confirm the plasma membrane localization of CNGC13, under its native promoter (**Figure 2F, G**). Using subcellular fractionation and western blot, we detected full length CNGC13-YFP protein expression exclusively in the plasma membrane fraction (**Figure 2H**), supporting the conclusion that CNGC13 is a plasma membrane-localized protein.

**Figure 2:**
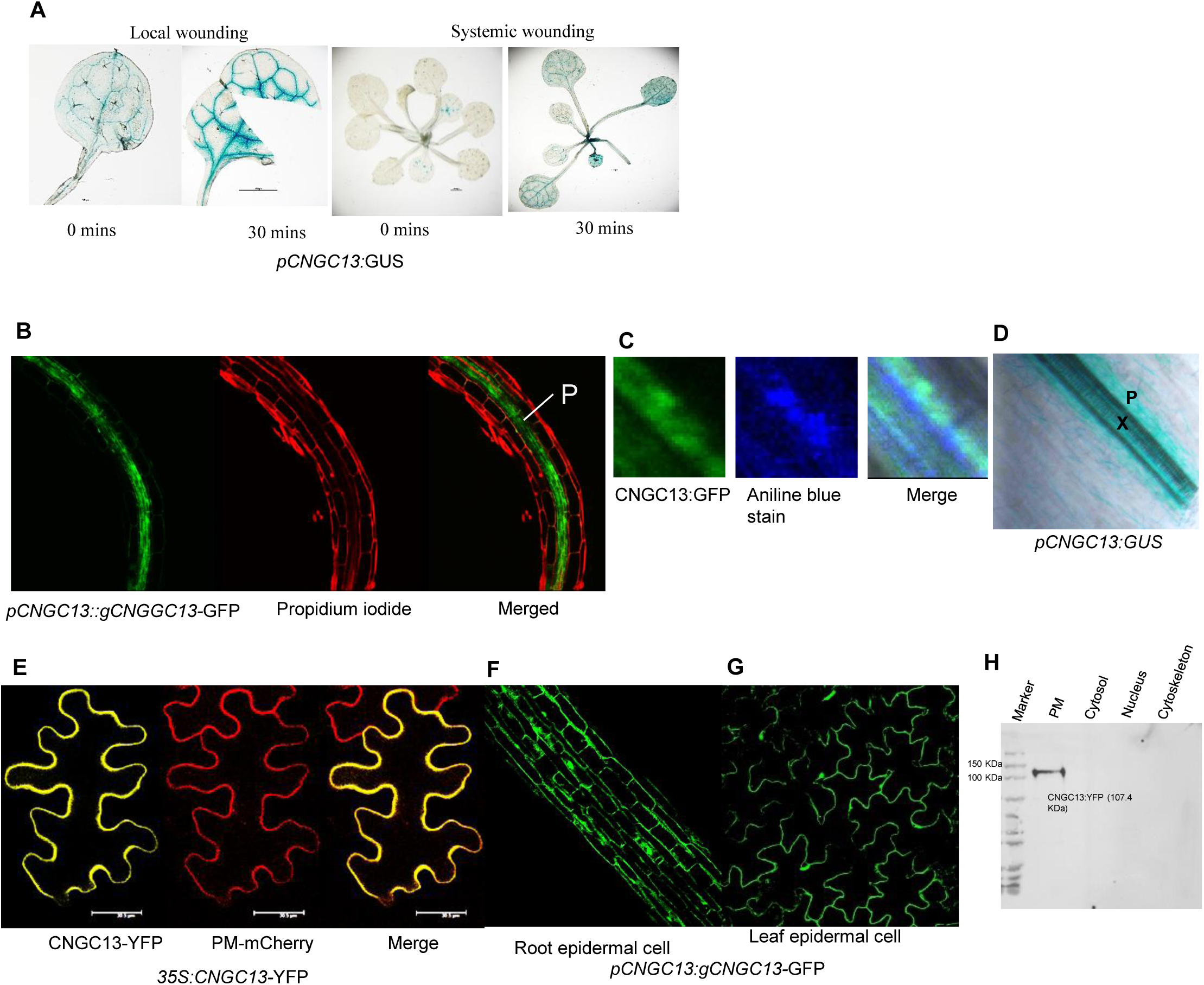
CNGC13 is expressed in vasculature and is localized at plasma membrane. (**A**) *CNGC13* promoter activity in leaf vasculature visualized by *CNGC13pro:GUS* fusion expressing transgenic plants; control or mechanically wounded, by cutting the midrib-Local wounding (left) and cutting petiole and imaging systemic leaves-Systemic wounding (right), after 30 min. After the treatments, leaf tissues/seedlings were collected and incubated in GUS staining solution for 16 hrs. (**B**) Confocal microscope images of *pCNGC13:gCNGGC13-*GFP plants in primary root, stained with propidium iodide (red) to stain the cell wall thus marking the cell boundaries. The sample was fixed, cleared, and then stained with aniline blue and propidium iodide (red). (**C**) Image shows that aniline blue stains a region of strong CNGC13-GFP accumulation in the phloem. Aniline blue stains callose and used as a marker for identification of sieve elements of phloem (**D**) *CNGC13pro:GUS* expression in longitudinal section of petiole vasculature from 2-week old *Arabidopsis* plant. P and X represents phloem and xylem respectively. (**E**) Visualization of CaMV35S*:CNGC13-YFP* fusion full-length protein in transiently transformed *N. benthamiana* plants using confocal laser scanning microscopy. Plasma membrane marker, *PM-mCherry* was co-infiltrated to facilitate CNGC13-YFP subcellular localization. Scale bar = 30.5 um. (**F, G**) *proCNGC13:gCNGC13-GFP* expression in root epidermal cell and leaf epidermal cell of 2-week-old *Arabidopsis* plant. (**H**) Subcellular fractionation to detect plasma membrane localized CNGC13-YFP after transient expression in *N. benthamiana* and *analysed* by western blot using anti-GFP antibodies.

### CNGC13 is a Ca^2+^-permeable channel

CNGCs are tetrameric ion channels and, although the Ca^2+^ permeability has been established for several members of this family (Dietrich et al., 2020), the ion conductance properties of CNGC13 remain uncharacterized. To address this, a codon optimized CNGC13 with YFP tag was expressed in *Xenopus laevis* oocytes. The oocytes that received the CNGC13-YFP cRNA injection displayed strong fluorescence at their edges, while controls did not exhibit any fluorescence. The fluorescence was confined to the edge of the oocyte, where a line scan indicates the fluorescence is restricted to the oocyte boundary (**Figure 3A**). Two Electron Voltage Clamp (TEVC) experiments were performed on CNGC13-YFP-expressing oocytes, by subjecting them to pulses ranging from −130 mV to +30 mV. No currents were seen in un-injected oocytes subjected to both cAMP and Ca^2+^ (**Figure 3B**). Large inward currents were observed in CNGC13-YFP expressing oocytes in the presence of external CaCl_2_ (15 mM) in the recording buffer and dibutyryl cAMP (300 µM), upon hyperpolarizing to <-90 mV (**Figure 3C**). *Xenopus* oocytes express Ca^2+^–activated chloride current that is activated upon elevation of cytosolic Ca^2+^. Treatment with the Ca^2+^–activated chloride current inhibitor, 4, 4’-diisothiocyanatostilbene-2, 2’-disulfonic acid disodium salt (DIDS; 300 µM), results in a reduction of the Ca^2+^–activated inward current (**Figure 3C**). We also tested the ion selectivity of the channel using monovalent cation K^+^, and near-background currents were observed upon its addition in the presence of cAMP (**Supplementary Figure 2A**). We also expressed full length CNGC13 in yeast *cch1/mid1* mutant to test its possible channel activity (Fischer et al., 1997). CNGC13 complements the growth-arrest phenotype of the *cch1/mid1* yeast mutant defective in mating pheromone-induced cytosolic Ca²⁺ signalling, indicating its Ca²⁺ permeability (**Supplementary Figure 2B**). These data establish that CNGC13 is a Ca^2+^ permeable channel.

**Figure 3:**
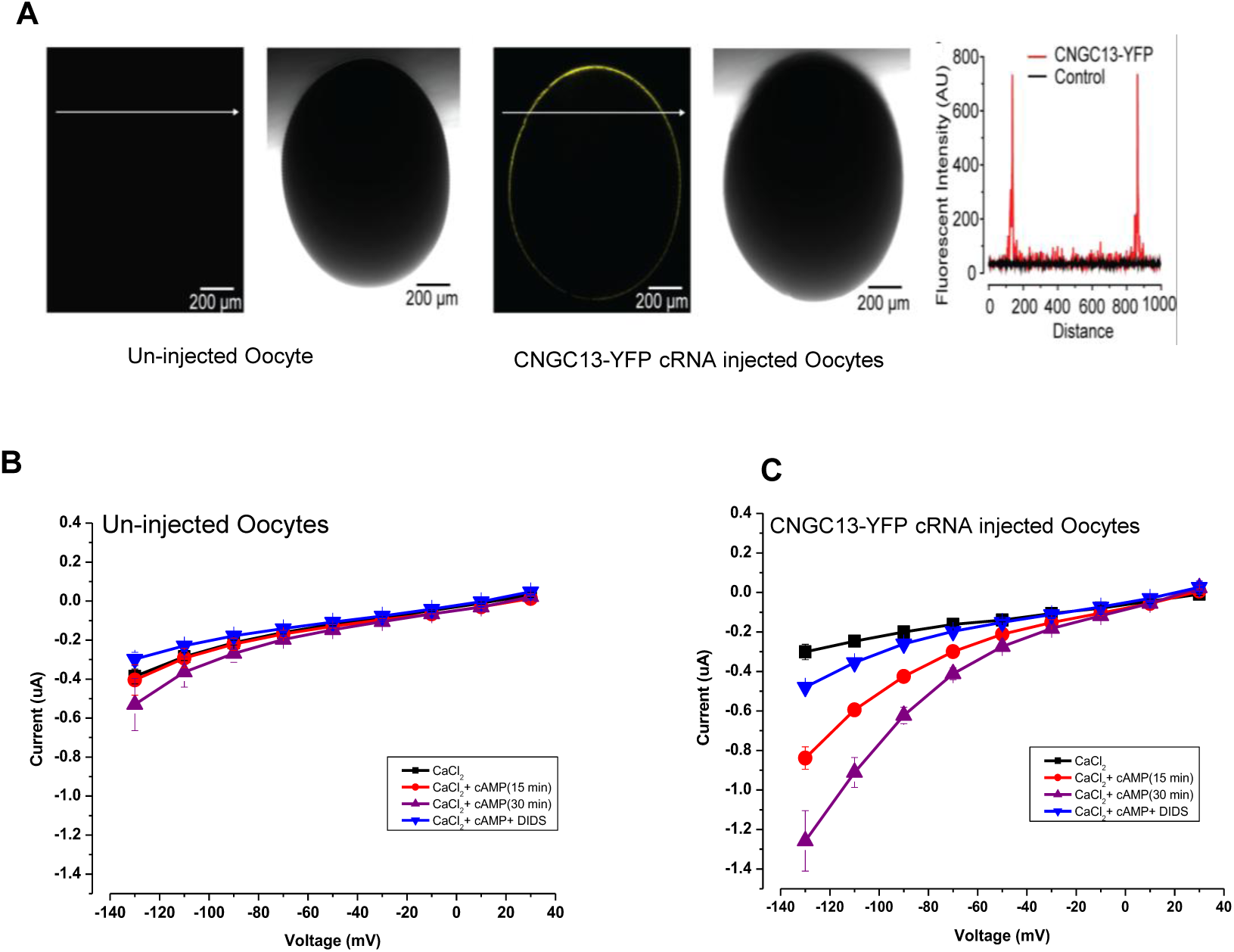
CNGC13 is a plasma membrane localized Ca^2+^ permeable channel. (A) Confocal imaging of CNGC13-YFP fusion protein expression in *Xenopus* oocyte. Fluorescence confocal images of an un-injected oocyte (left panel), and its bright field image (right panel), fluorescence confocal image of CNGC13-YFP cRNA injected (left panel), and its bright field image (right panel). Line scans of fluorescence intensities along with lines drawn in (i; left panel) and (ii; left panel). Distance (x-axis) is from the start of the arrow. Red-CNGC13-YFP injected oocyte; Black-un-injected oocyte. Scale bar: 200 μm. (B, C) TEVC recordings for un-injected and CNGC13-YFP expressing oocytes. Currents were recorded on a un-injected and CNGC13 channel cRNA injected oocyte in CaCl_2_ (15 mM) recording buffer, before and after the addition of dibutyryl cAMP (300 µM) for 15 min and 26 min, which is followed by addition of DIDS (300 µM). All additions were done sequentially and no washout was performed for the recordings. In panel (B) Plots of current values recorded from un-injected oocytes and (C) Plots of current values recorded from CNGC13-YFP cRNA injected oocytes, in CaCl_2_ (15 mM) recording buffer before and after the addition of 300 µM dibutyryl cAMP (post 15 min and 26 min) followed by addition of DIDS (300 µM). Each data point is an average obtained from n = 4, and the error bars correspond to standard error values.

### CNGC13 is crucial for Ca^2+^ signal propagation to distal leaves upon wounding

Given the vascular localization and wound-inducible transcriptional activation of *CNGC13*, we hypothesized that this channel contributes to wound-induced cytosolic Ca^2+^ wave propagation. To test this, we transformed *cngc13-1* and *cngc13-3* mutants with GCaMP3 Ca^2+^ indicator (Tian et al., 2009), and selected *cngc13-1*_GCaMP3_ and *cngc13-3*_GCaMP3_ stable lines with comparable basal fluorescence to wild type (WT) (**Supplementary** Fig. 3A**, 3B**). In WT plants, petiole cutting triggered a rapid cytosolic Ca^2+^ increase in the wounded local leaf (LL), which propagated via the vasculature to systemic leaves. On cutting petiole at 30 s, Ca^2+^ signals appeared in the LL petiole at ∼45 s, reached the systemic petiole by ∼75 s, and spread to the systemic leaf lamina by ∼90 s, persisting until ∼150 s before declining **(Figure: 4A, Supplementary Movie S1)**. In contrast, both *cngc13* mutants exhibited a delayed and attenuated cytosolic Ca^2+^ increase. While the initial Ca²⁺ rise in the LL persisted and reached the LL petiole by ∼45 s, signal propagation beyond this point was markedly impaired. In the majority of mutants (12/16), the LL petiole signal remained static until ∼90 s, with weak Ca^2+^ signals emerging in the systemic petiole only after ∼105 s and reaching the lamina at ∼150 s **(Figure: 4B, 4C, Supplementary Figure 4, Supplementary Movie S1-S3)**. These experiments were conducted in 2-week-old plants, as GCaMP3 silencing was observed in mature *cngc13* mutant plants at 4–5 weeks of age. Quantification of ΔF/F₀ over time revealed significantly reduced and delayed systemic Ca^2+^ responses in both *cngc13* mutants compared to WT (**Figure 5A-C**). Additionally, propagation velocities measured across defined regions of interest (ROIs) were consistently lower in *cngc13* lines (**Figure 5D**). These findings suggest that CNGC13 is essential for the efficient transmission of wound-induced Ca^2+^ signals in Arabidopsis.

**Figure 4:**
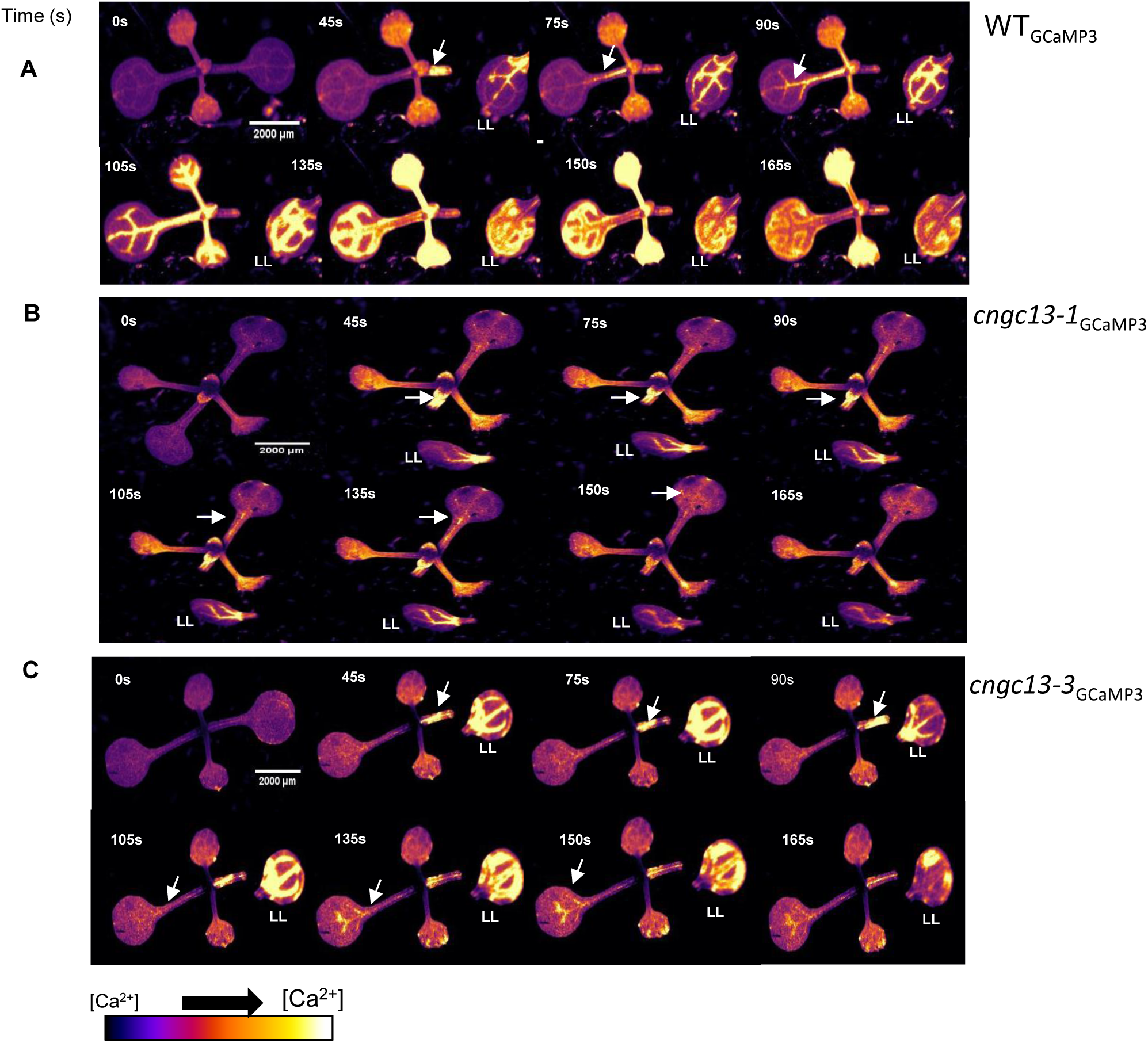
CNGC13 is involved in generation of systemic Ca^2+^ elevation upon wounding. **(A), (B)** and **(C)** Representative stereomicroscope images showing fluorescence changes upon wounding (cutting petiole) at different time intervals in wild type, *cngc13-1* and *cngc13-3* seedlings in GCaMP3 background (T3 generation). Cutting leaf 1 petiole in a 12-15-day-old plant caused a systemic [Ca**^2+^**]_cyt_ increase that propagated throughout the rosette. n=8 independent experiments. Scale bar = 2000 µm. White arrows represent the path of Ca**^2+^** signal movement from petiole of cut leaf to petiole of systemic leaf to whole leaf. LL= Local Leaf (The leaf whose petiole was cut)

**Figure 5:**
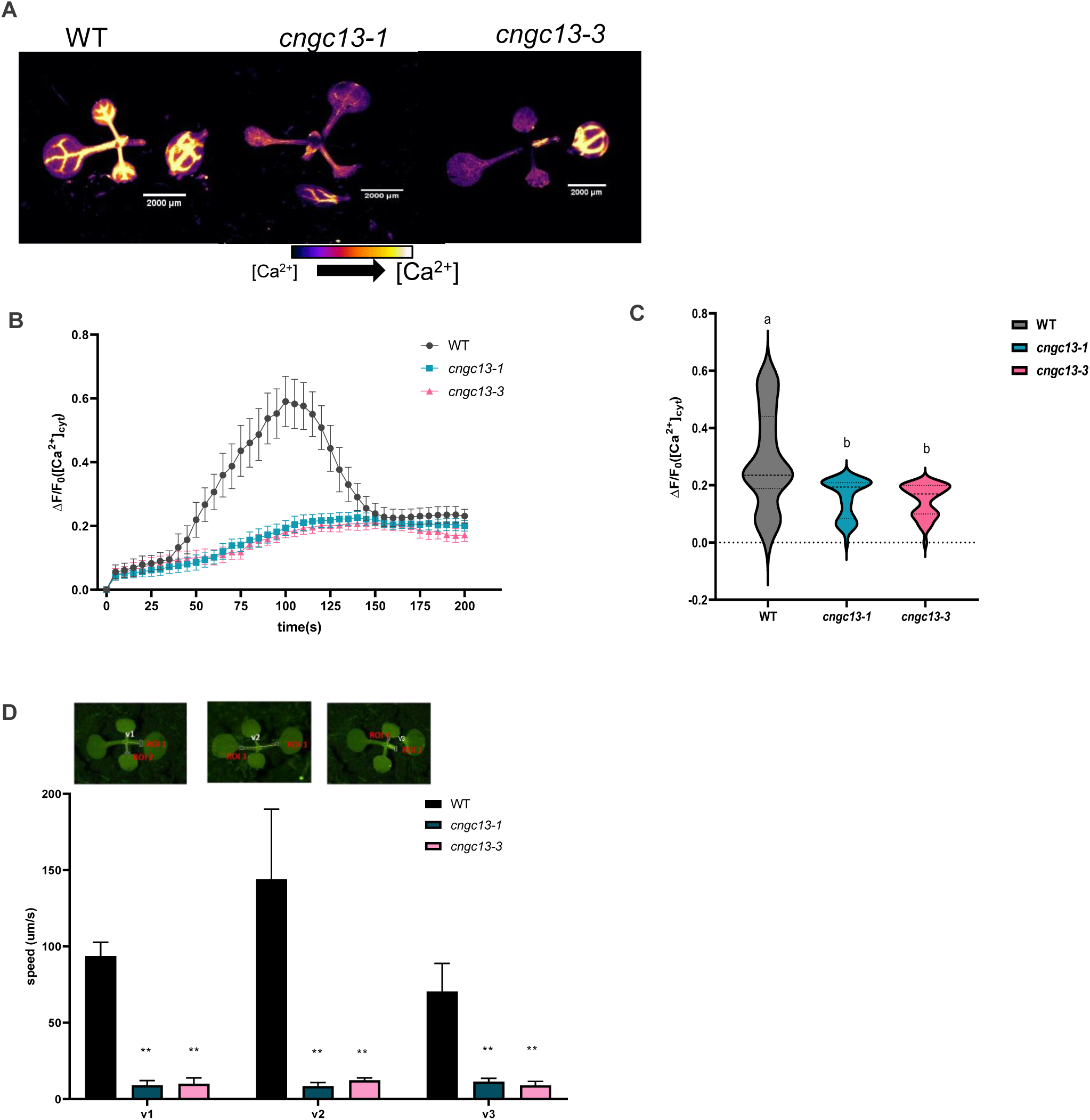
CNGC13 regulates systemic Ca^2+^ elevation upon wounding. (**A**) Representative stereomicroscope images showing fluorescence changes upon wounding at 105 s interval in WT, *cngc13-1* and *cngc13-3.* (**B**) Mean of normalized GCaMP3 fluorescence intensity ΔF/F_0_ and (**C**) Maximum GCaMP3 fluorescence intensity ΔF/F_0_ in systemic leaf of WT (black) *cngc13-1* (blue) and *cngc13-3* (pink) at 5 s intervals after wounding the leaf with fine scissors. F_0_, average fluorescence intensity before wounding (baseline); ΔF, difference between measured fluorescence and baseline fluorescence. The entire leaf area was selected for calculating ΔF/F_0_. One-way ANOVA: *P* < 0.05. Error bars represent the Mean ± SE of the mean (n = 8). (**D**) Speed of [Ca**^2+^**]_cyt_ wave propagation from site of cutting to systemic leaf at four leaf stage. Diagram showing *Arabidopsis* leaves and the regions of interest (ROI) used to analyze the [Ca**^2+^**]_cyt_ changes and speed (v1-v3) from the long-distance transmission of [Ca**^2+^**]_cyt_ increases. v1 is between ROI1 and ROI2 on the petiole of wounded leaf; v2 is between ROI1 and ROI3 on the petiole of wounded leaf; v3 is between ROI1 and ROI4 on the petiole of wounded leaf. Speed (µm/s) of the long-distance elevation in cytosolic Ca**^2+^** monitored using the Ca**^2+^** indicator GCaMP3 expressed in WT and *cngc13* mutant lines. An asterisk denotes statistical differences (*, *P* < 0.05) from WT data. The increase in signal used to calculate speed was defined as an increase to above 2 SD of the pre-stimulation levels. Error bars represent Mean ± SE, n = 4-5 replicates.

### CNGC13 mediated leaf-to-leaf Ca^2+^ propagation depends on ricca factor thioglucosidase and aliphatic glucosinolate aglucones

Leaf-to-leaf wound-response signalling in plants involves myrosinase/thioglucosidase (TGG) proteins as the principal mobile component and as they migrate through the vasculature, these TGGs catalyze the localized hydrolysis of glucosinolates (GSs). Metabolite profiling revealed rapid wound-induced breakdown of aliphatic glucosinolates in primary veins producing aglucone intermediates that initiate sustained membrane depolarization and cytosolic Ca^2+^ transients in systemic leaves (Gao *et al*., 2023). Glutamate, another known mobile elicitor, serves as a ligand for the phloem-localized glutamate receptor-like (GLR) channel GLR3.3 (Alfieri et al., 2020; Grenzi et al., 2023), which, together with GLR3.6, mediates long-distance electrical and Ca²[ signalling (Nguyen et al., 2018; Toyota et al., 2018). To assess the involvement of these pathways in *cngc13* mutants, we analyzed the expression of *GLR3.3* and *GLR3.6* in WT and *cngc13* plants by cutting petiole of leaf 8 and analyzing gene expression in the vascularly connected leaf 13. GLRs showed activation at 30 and 60 minutes, respectively, with no differential expression between genotypes (**Figure 6A, 6B)**. In contrast, *TGG* expression peaked at 30 minutes in WT but was undetectable in the *cngc13-1* background, suggesting a potential link between CNGC13 function and TGG-mediated signal propagation (**Figure 6C, 6D)**. To test the hypothesis that CNGC13 is required for systemic wound signalling by enabling TGG-dependent GS breakdown and accumulation of bioactive isothiocyanates (ITCs) in distal tissues, we performed LC-MS/MS-based metabolite profiling. Following crush wounding of leaf 8, we harvested the midrib of leaf 13, 2 minutes post-injury and quantified both intact aliphatic GSs and their ITC breakdown products, which serve as proxies for short-lived aglucones. In WT, levels of Glucoraphanin-ITC and Glucoalyssin-ITC were significantly elevated in wounded veins compared to unwounded controls. In contrast, no such increase was observed in both *cngc13* mutant veins upon wounding (**Figure 7A, 7B)**. Levels of intact GSs remained unchanged between wounded and unwounded WT and *cngc13* in the veins (**Supplementary** Fig. 5), indicating that the defect lies in GS hydrolysis rather than transport or biosynthesis in the veins. These findings establish CNGC13 as a critical regulator of systemic wound signalling, acting upstream of TGG-dependent aliphatic glucosinolate hydrolysis to generate aglucones that drive Ca^2+^ elevation in distal leaves.

**Figure 6:**
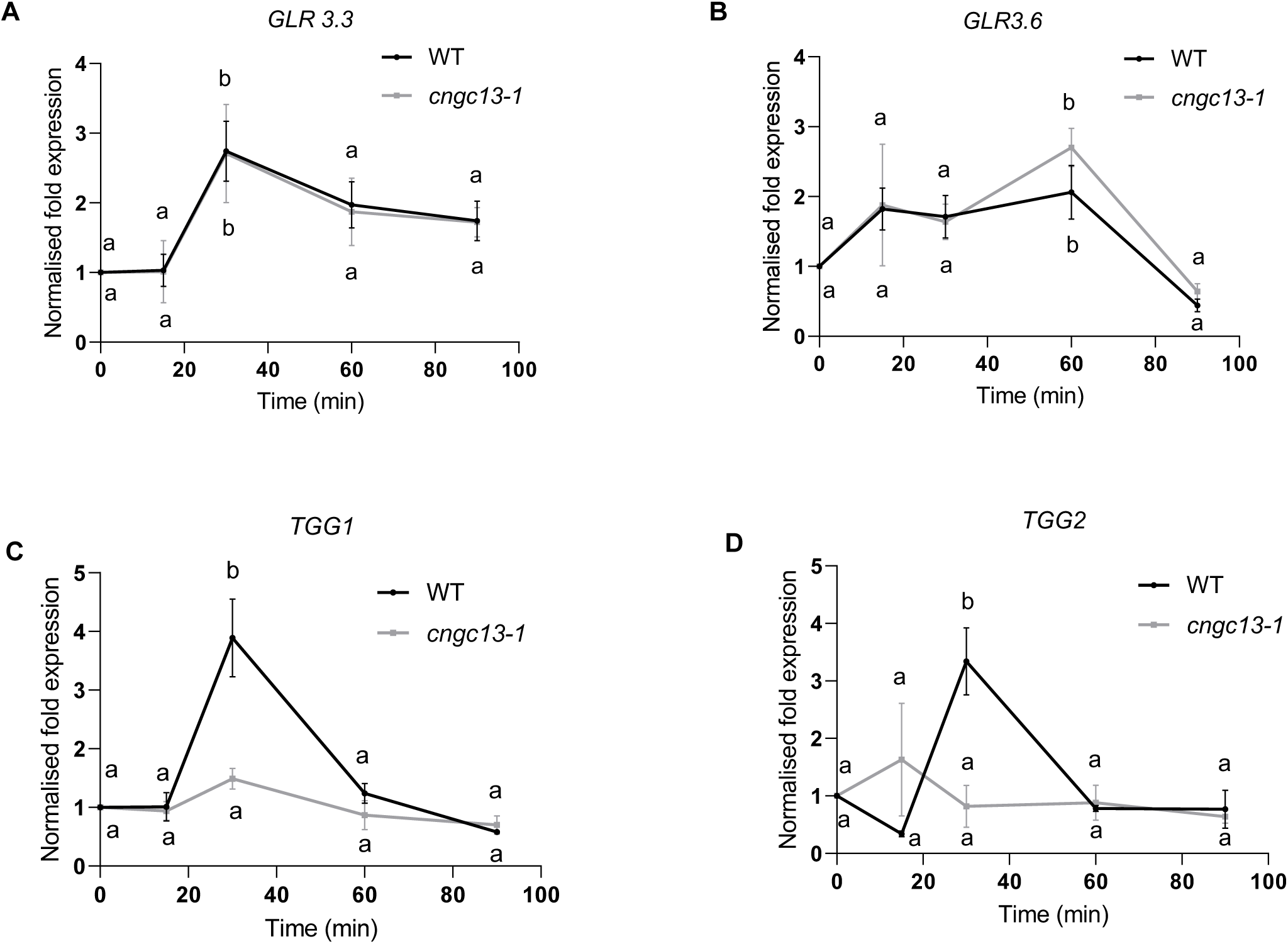
Transcript accumulation of systemically expressed genes in WT and *cngc13* upon wounding. (A) *GLR3.3* and (B) *GLR3.6* (C) *TGG1* (D) *TGG2* transcript induction in leaf 13 of WT and *cngc13-1* at 0, 15, 30, 60 and 90 min after cutting the petiole of leaf 8. The graphs show fold change (Mean ± SE) of mRNA level relative to untreated control. The figure shows data from the means of three independent experiments with n= 6. Different letters indicate statistically significant differences among treatments, calculated by one-way ANOVA

**Figure 7:**
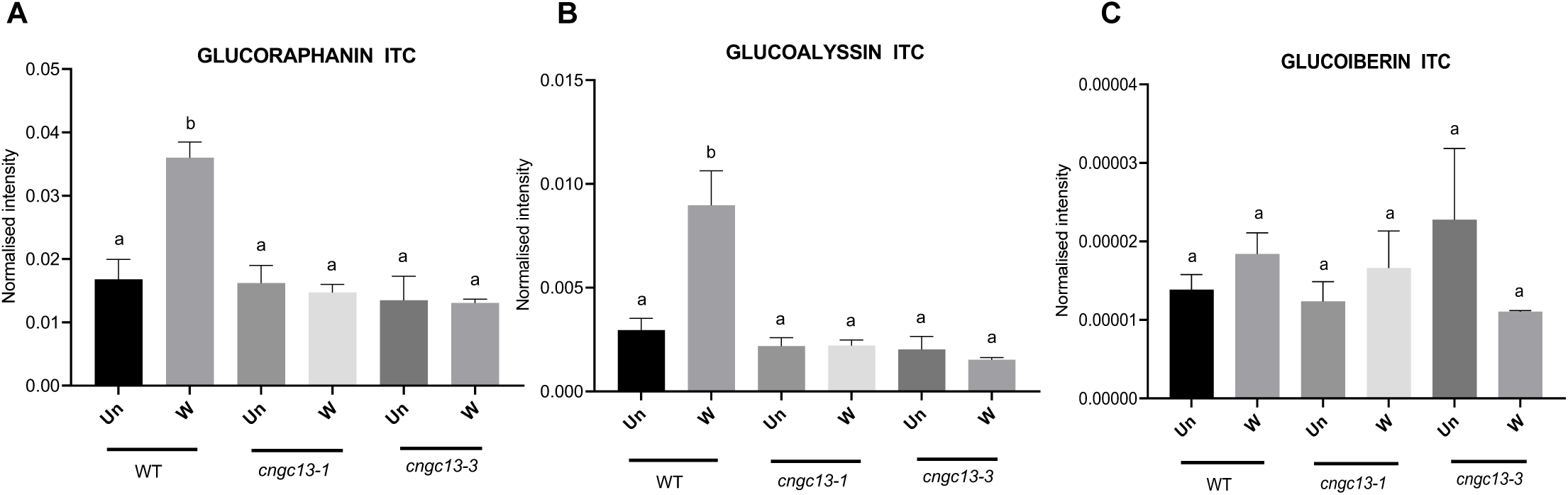
Systemic Ca^2+^ propagation depends on ricca factor and short-lived glucosinolate aglucone elicitors. Quantification of isothiocyanates (ITCs) from aliphatic glucosinolates in midrib (**A**) Glucoraphanin-ITC, (**B**) Glucoalyssin-ITC, (**C**) Glucoiberin-ITC in WT, *cngc13-1* and *cngc13-3* mutant lines upon control or unwounded (Un) and wound (W) treatment. ITCs were quantified after crush wounding leaf 8 and collecting midrib of leaf 13 after 2 min of treatment. The figure shows Mean + SE values data from three independent experiments with n = 3 (each replicate is a pool of 5-6 leaf midrib collected from different plants; 3*6 = 18 plants). Different letters indicate statistically significant differences among different genotypes after treatment, calculated by one-way ANOVA followed by Tukey’s test (*P* ≤ 0.05). WT, wild type.

### Systemic induction of jasmonate biosynthesis is hampered in *cngc13* mutants

Wounding induces rapid hydraulic and electrical changes that activate Ca^2+^ influx, which in turn initiates jasmonate biosynthesis in vascularly connected systemic leaves (Farmer et al., 2014; 2020). Notably, *CNGC13* expression is upregulated as early as 15 minutes post-wounding and peaks at 60 minutes (**Figure 1A**), temporally coinciding with the onset of jasmonic acid (JA) biosynthesis (Vadassery et al., 2012). To determine whether CNGC13 influences wound-induced JA production, we quantified jasmonate levels in the systemic leaf (L13) of WT and *cngc13* mutants following wounding of a leaf 8 (L8, local leaf), after 15 minutes. In WT plants, wounding L8 led to significant increases in both JA and JA-isoleucine (JA-Ile) levels in L13. By contrast, systemic accumulation of both JA and JA-Ile in L13 was highly reduced in both *cngc13* alleles upon wounding (**Figure 8A and 8B**). Levels of other hormone like ABA/SA levels were unaltered in systemic leaves (**Supplementary** Fig. 6A**, 6B**). LOX6 activity drives the rapid systemic synthesis of JA and JA-Ile, while JAZ10 serves as a key marker for distal jasmonate biosynthesis (Chauvin et al., 2013; Mousavi et al., 2013). Consistent with this, transcript levels of the jasmonate-responsive marker gene *JAZ10* and *LOX6* were significantly attenuated in L13 of wounded *cngc13* plants, resembling the response observed in *glr* mutants (Mousavi et al., 2013) (**Figure 8C and 8D**). Importantly, no significant difference in jasmonate levels were observed between WT and *cngc13* plants in the local leaf upon both wounding (**Supplementary Fig. 7A, 7B**) or herbivore feeding (**Supplementary Fig. 7C, 7D**). These results indicate that *CNGC13* is specifically required for the systemic, but not local, induction of jasmonates and associated immune responses following wounding.

**Figure 8:**
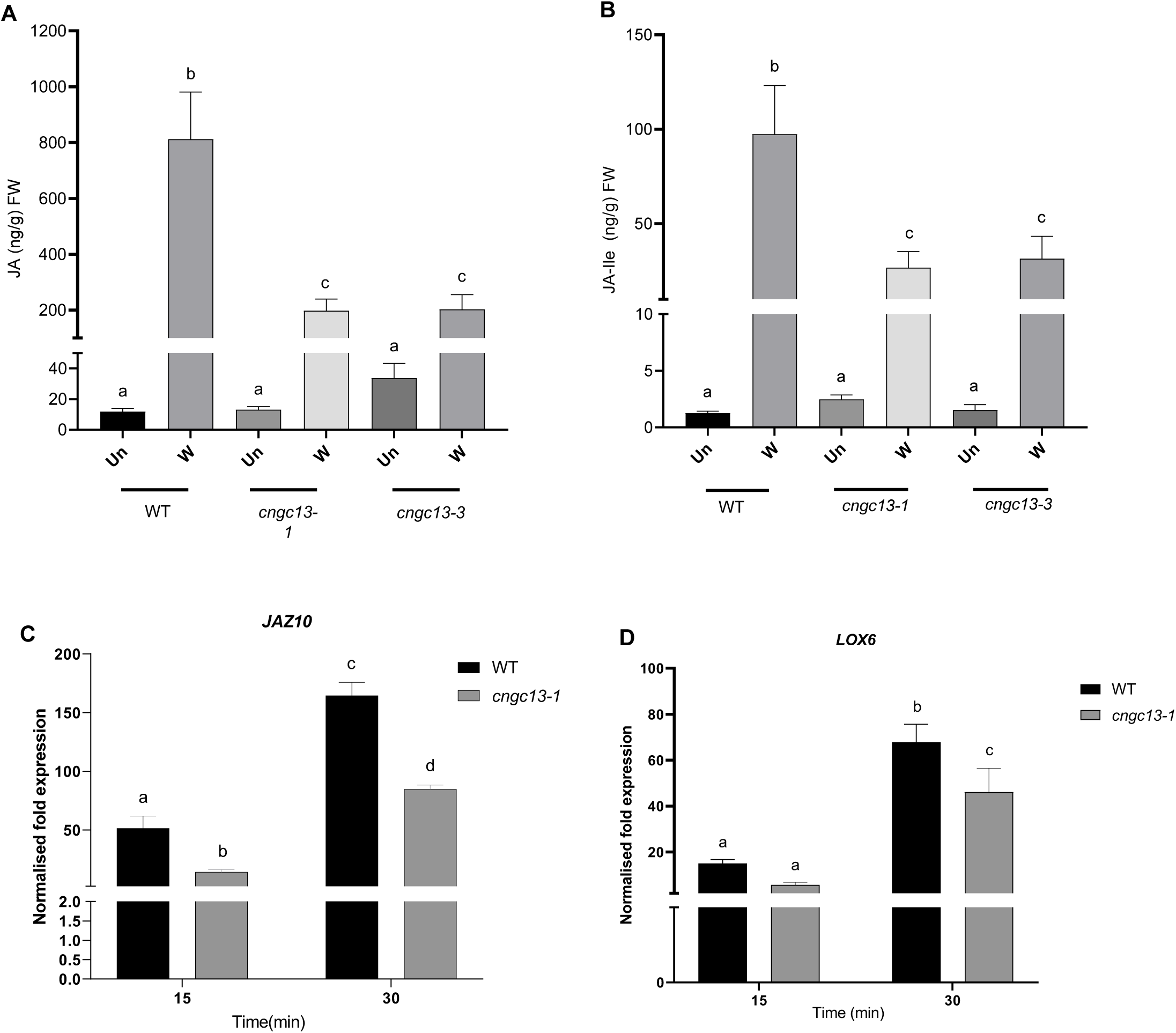
CNGC13 is involved in systemic jasmonate accumulation on wounding. (**A**) JA and **(B)** JA-Ile phytohormone elevation upon wound treatment in WT, *cngc13-1* and *cngc13-3* plants. Mean + SE values are shown for JA and its bioactive form -JA-Ile in *Arabidopsis* WT, *cngc13-1 cngc13-3* (dark grey) plants in unwounded (Un) and systemic leaf wounded (W) conditions. The phytohormone levels were measured from systemic leaf 13, collected 15 min after cutting the leaf 8 petiole. The figure shows data from three independent experiments with n = 6 (each replicate is a pool of 3 leaves collected from 3 different plants; 3*6 = 18). Different letters indicate statistically significant differences among treatments, calculated by one-way ANOVA followed by Tukey’s test (*P* ≤ 0.05). WT, wild type. **(C)** *JAZ10* (**D**) *LOX6* transcript induction in systemic leaf 13 of WT and *cngc13-1* at 15min and 30 min, after cutting the petiole of leaf 8. The graphs show fold change (Mean ± SE) of mRNA level relative to untreated control. The figure shows data from the means of three independent experiments with n=6. Different letters indicate statistically significant differences among treatments, calculated by two-way ANOVA followed by Tukey’s test (*P* ≤ 0.05). WT, wild type.

### CNGC13 is required for *At*Pep-induced cytosolic Ca²**^⁺^** and ROS signalling

Damage-associated molecular patterns (DAMPs) such as *At*Peps are recognized by plants upon herbivory (Klauser, 2015). Analysis of previously published RNA-seq data revealed that *CNGC13* is induced after 2 h of *At*Pep treatment (Ross et al., 2014). Genevestigator-based co-expression analysis further showed that *CNGC13* is upregulated in response to *At*Peps in a BAK1-dependent manner (**Supplementary** Fig 8A). Consistent with this, we found that *CNGC13* transcript levels increased rapidly upon *At*Pep1 and *At*Pep2 treatment, peaked at 60 minutes, and remained elevated for up to 6 h (**Figure 9A**). To study its spatial activation, we examined CNGC13 promoter activity in *proCNGC13::GUS* lines. Upon *At*Pep1 and *At*Pep2 treatment, reporter activity was restricted to the leaves and absent in stems and roots (**Supplementary** Fig. 8B). To test whether CNGC13 contributes to *At*Pep-triggered Ca²⁺ signalling, we introduced a cytosol-targeted aequorin reporter (Knight et al., 1997) into *cngc13-1* and *cngc13-3* mutant backgrounds. In WT seedlings, both *At*Pep1 and *At*Pep2 triggered rapid and sustained cytosolic Ca²⁺ elevations within 1 min of treatment. In contrast, *At*Pep1 and *At*Pep2 induced Ca^2+^ signals were reduced in both *cngc13* mutant alleles (**Figure 9B, 9C**). Since CNGC13 is involved in systemic response, we investigated whether *At*Peps induces a systemic signal and we could not detect any systemic signal even in WT-GCaMP3 reporters. This could also be that like glutamate (applied in mM concentration), liquid application of *At*Peps is not strong enough to induce systemic signalling (Grenzi et al., 2023). As *At*Peps also activate Reactive Oxygen Species (ROS) production as part of PTI (Klauser et al., 2014), we next tested whether CNGC13 affects this response. *At*Pep2 failed to induce detectable ROS in either WT or *cngc13* mutants. However, *At*Pep1 triggered a robust and prolonged ROS burst in WT, which was significantly attenuated in both *cngc13-1* and *cngc13-3* alleles (**Figure 9D**). Expression of ROS-related genes *RBOHD*, *WRKY33*, and *RRTF1* was also significantly lower in the *cngc13* mutants compared to WT following *At*Pep1 treatment (**Supplementary** Fig 8C). Together, these findings identify CNGC13 as a key component of the *At*Pep signalling cascade that mediates early Ca^2+^ influx and ROS-based defense activation to downstream defense responses in Arabidopsis in local leaf.

**Figure 9:**
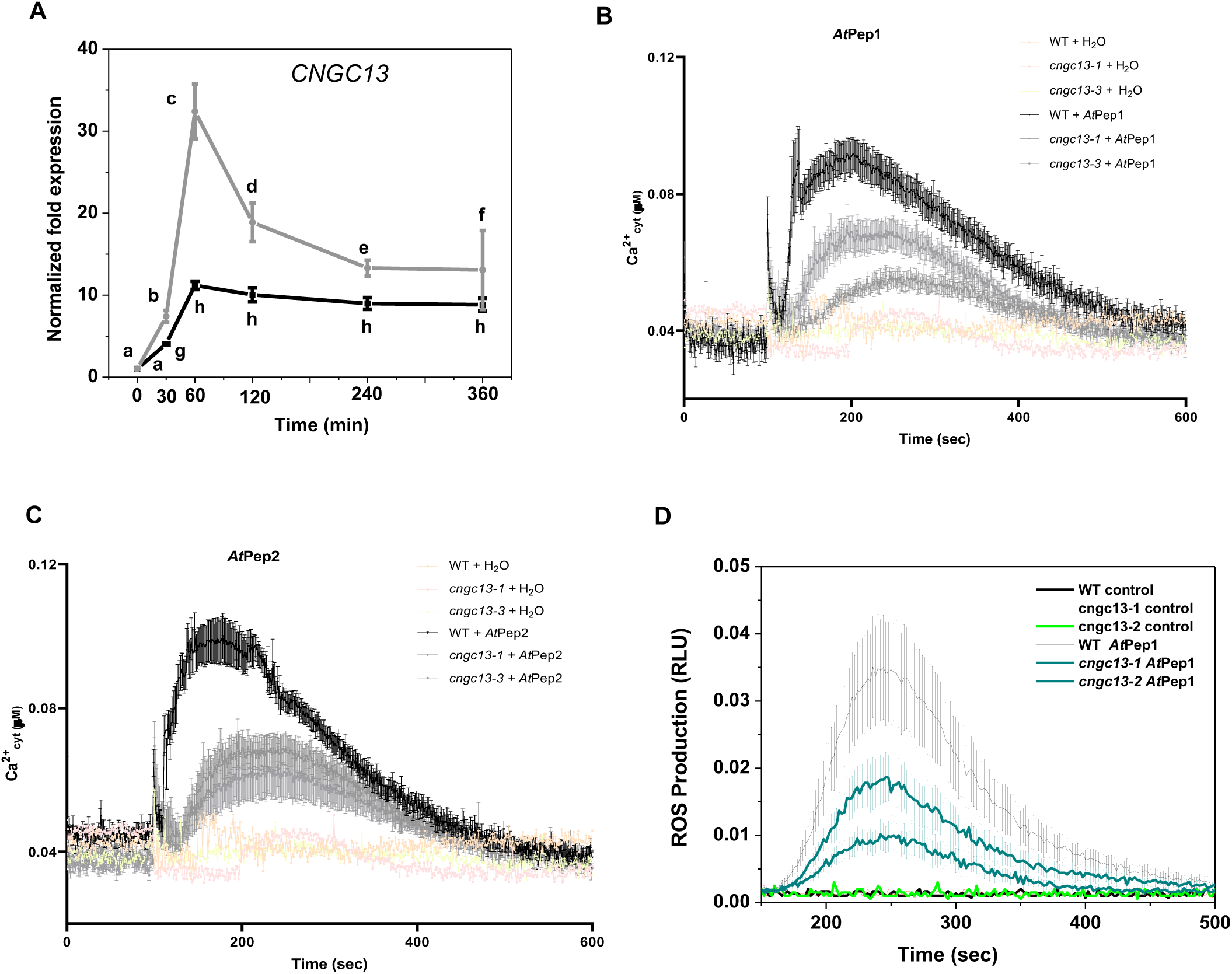
CNGC13 is involved in *At*Pep induced Ca^2+^ elevation and ROS burst. **(A)** *CNGC13* transcript accumulation in 14 d old *Arabidopsis* seedling treated with 10 nM of *At*Pep1 (black) and *At*Pep2 (grey) for different time points. The graphs show fold change (Mean ± SE) of mRNA level relative to untreated control. The figure shows data from the means of three independent experiments with n = 4. **(B)** 100 nM *At*Pep1 and **(C)** *At*Pep2 induced cytosolic Ca**^2+^** elevation (Ca**^2+^**_cyt_) in 10-day–old WT aequorin-*pMAQ2* (green) and *cngc13-1-pMAQ2* mutant (blue) seedlings. (Ca**^2+^**_cyt_) elevation response was captured with 300 ms integration time and 1 sec interval time. Remaining Ca**^2+^** was discharged using 2 M and 20% CaCl_2_. Water was used as control (black and red). Values are mean ± SE (n = 8). **(D)** Mean ± SE of ROS production in WT (black), *cngc13-1* (cyan) and *cngc13-2* (cyan) mutant plants upon 1 µM *At*Pep1 treatment (n = 12). *Arabidopsis* seedlings were grown for 12 d in liquid MS and treated with 1 µM *At*Pep1 or water as control.

### Loss of CNGC13 reduces aliphatic glucosinolate accumulation and enhances herbivory

To determine whether CNGC13 contributes to downstream secondary metabolite defense against herbivory, we analyzed whole-plant glucosinolate (GS) levels in wild-type and *cngc13* mutants under both basal and insect-challenged conditions. Under control conditions, *cngc13* mutants exhibited lower total GS levels compared to WT plants in the whole plant (**Figure 10A**). This reduction was attributed specifically to a decrease in aliphatic GSs, whereas indolic GS levels remained unchanged (**Figure 10A**). It is important to note that this difference is at the whole plant level only and not specifically in leaf veins. We further assessed GS accumulation after 7 days of *S. litura* herbivory. While WT plants showed an increase in total GS content on herbivory, *cngc13* mutants maintained significantly lower GS levels after 7 days of *S. litura* feeding (**Figure 10B**). These results indicate that CNGC13 is required for proper aliphatic GS accumulation under both basal and herbivore-induced conditions in whole plants, and that its loss compromises secondary metabolite defense, potentially contributing to enhanced herbivory.

**Figure 10:**
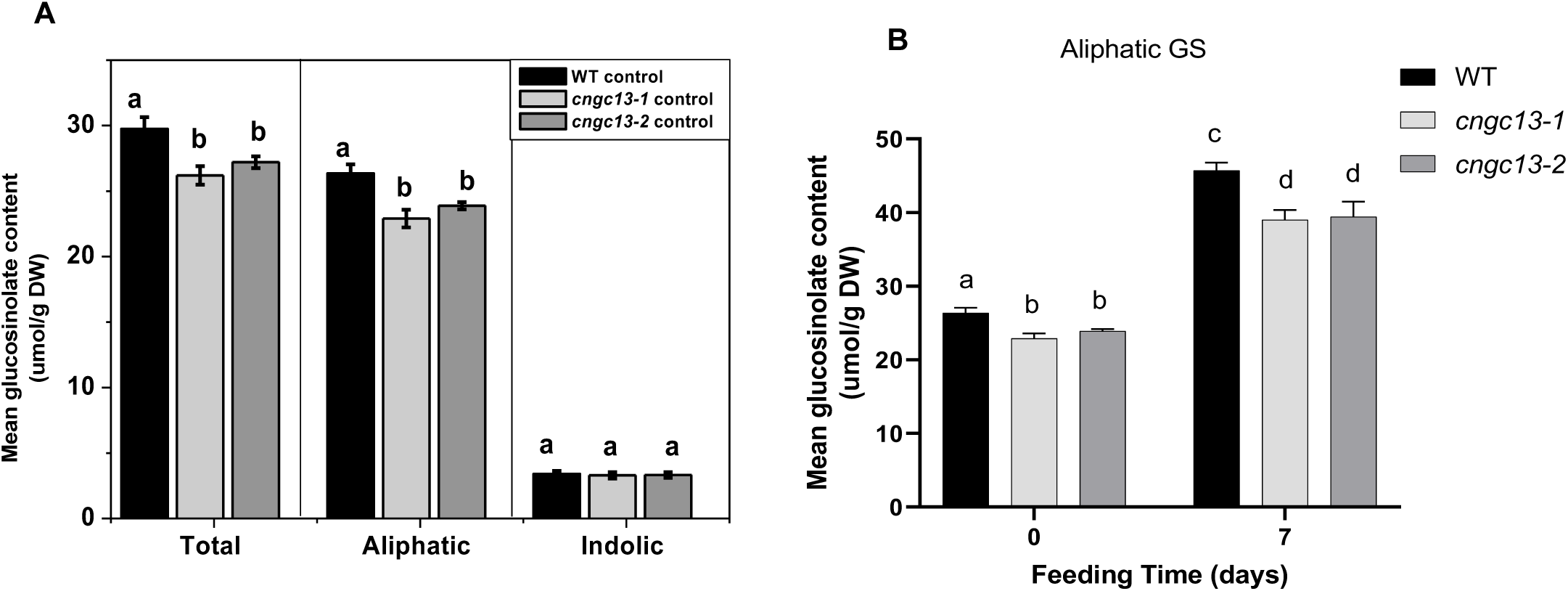
CNGC13 affects basal level and herbivory induced glucosinolate levels. (**A**) Mean ± SE (n=5) total GS levels in whole rosette of 5-week-old *Arabidopsis* plants at basal level in WT (black), *cngc13-1* (light grey) and *cngc13*-2 (grey) plants. Statistically significant differences between plants were analyzed by one-way ANOVA and Tukey HSD *P* ≤ 0.05. (**B**) Mean ± SE (n=5) of aliphatic GS levels in rosette of 5-week-old *Arabidopsis* plants at basal and after 7 d feeding with fourth instar *Spodoptera* larvae on WT, *cngc13-1*, and *cngc13*-2 plants. Statistically significant differences between plants were analyzed by one-way ANOVA and Tukey Test, *P* ≤ 0.05.

### CNGC13 and CNGC19 work together to provide plant defense against herbivory

We have previously shown the role of CNGC19 as a positive regulator of *S. litura* herbivory (Meena et al., 2019). In Arabidopsis pollen tubes, it has been proposed that Ca^2+^-permeable channels such as GLRs can partially compensate for each other’s loss by upregulating the expression of redundant members (Wudick et al., 2018). To assess whether cyclic nucleotide-gated channels (CNGCs) might exhibit similar compensatory dynamics, we analyzed the expression of *CNGC13* in the *cngc19* mutant background and CNGC19 in the *cngc13* background during simulated herbivory. At 60 minutes, both W+W and W+OS treatments triggered significantly higher *CNGC13* expression in *cngc19-2* compared to WT plants (**Figure 11A**). Conversely, CNGC19 expression in *cngc13-1* mutants was higher than in WT plants upon W+W and W+OS treatments (**Supplementary** Fig. 9A). These results suggest a potential compensatory mechanism between CNGC13 and CNGC19. Given that both channels positively regulate defense responses to insect herbivory, we hypothesized that CNGC13 and CNGC19 might act cooperatively. To test this, we generated a *cngc13 cngc19* double mutant and performed insect bioassays. *S. litura* larvae feeding on the double mutant gained significantly more weight compared to those on WT, *cngc13*, or *cngc19* single mutants (**Figure 11B, 11C**), suggesting enhanced susceptibility. In line with these observations, the double mutant accumulated lower levels of aliphatic glucosinolates upon prolonged herbivory (**Figure 11D**). CNGCs are known to form functional homo- or hetero-oligomeric channels (Dietrich et al., 2020). To investigate whether CNGC13 and CNGC19 form heteromeric channels, we tested their interaction using Y2H. CNGC13 does not interact with CNGC19, nor does it form homodimers via its N- or C-terminal domains (**Figure 11E; Supplementary Fig. 9B**). These findings indicate that while CNGC13 and CNGC19 contribute cooperatively to herbivory resistance, they likely function through independent channel complexes rather than as direct physical partners.

**Figure 11:**
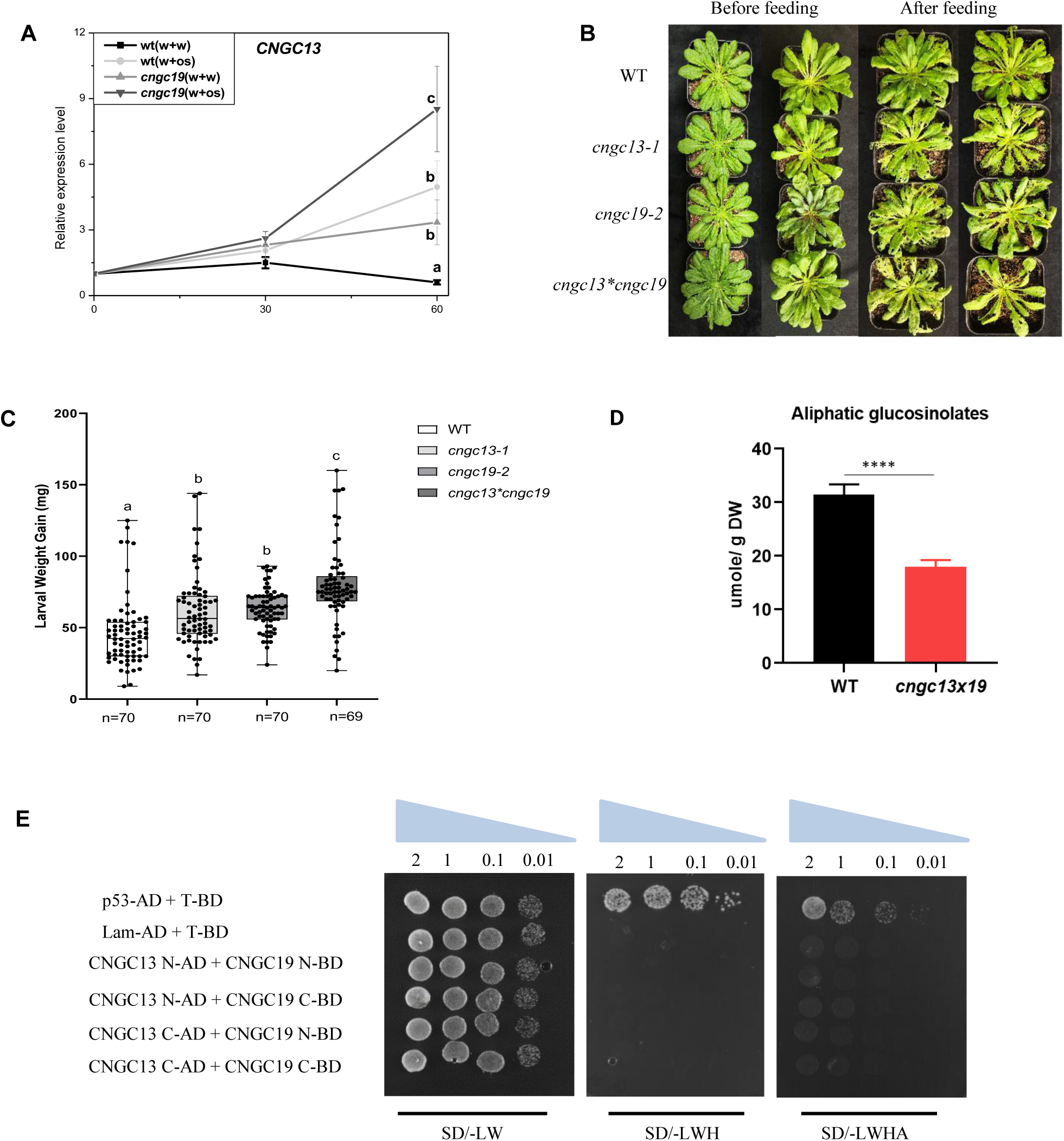
CNGC13 and CNGC19 have additive functions. **(A)** *CNGC13* transcript levels in 6-week-old *Arabidopsis* WT or *cngc19* mutant leaves after 30 and 60 min of simulated herbivory. Untreated plants were used as control for calculation. Plotted values are the mean ± SE (n = 4). Different letter indicates statistical significance difference among samples, calculated by one-way ANOVA (*P* ≤ 0.05). **(B)** Phenotype and leaf area eaten upon feeding *S. litura* larvae on WT, *cngc19-1*, *cngc13-1*, *cngc13*cngc19* for 8 d. **(C)** Larval weight after feeding on WT, *cngc19-1*, *cngc13-1*, *cngc13*cngc19* for 8 d. For all experiments *S. litura* 2^nd^ instar larvae growing in light for 3 d after hatching, were pre-weighed and three larvae were placed on plants of each genotype. The larval weight (mean ± SE) was measured after 7 d of feeding. The total number of larvae weighed (n) is indicated in the bars. Statistically significant differences between genotypes after feeding of the larvae were analyzed by one-way ANOVA using SNK test, *P*=<0.001. **(D)** Mean ± SE (n = 5) of aliphatic GS levels in rosette of 5-week-old *Arabidopsis* plants after 7 d feeding with fourth instar *Spodoptera* larvae on WT and *cngc13*cngc19* plants. Statistically significant differences between plants were analyzed by t-test; *p* ≤ 0.001. **(E)** Y2H interaction study of CNGC13 and CNGC19. Y2H gold yeast cell co-transformed with pGBKT7-CNGC13 N and C terminal and pGADT7-CNGC19 N and C terminal. The co-transformed constructs ten µL from a 10-time dilution series were spotted on control non-selective DDO and selective SD-Leu/-Trp/-His (TDO) and SD-Leu/-Trp/-His/-Ade (QDO) medium. Diminution of cell density in the dilution series is indicated by narrowing triangles. Y2H assay was performed four independent times. Each time similar observation was obtained, and the representative image is presented.

### CNGC13 interacts with pattern recognition receptor, PEPR2 on the plasma membrane

CNGC13, like other CNGCs, is localized to the plasma membrane, positioning it to participate in signalling events initiated by membrane-bound receptor kinases. Given that both CNGC13 and the *At*Pep receptors, PEPR1 and PEPR2 are plasma membrane-localized, and that CNGC13 contributes to *At*Pep-mediated responses, while PEPRs are known regulators of systemic immunity (Ross et al., 2014), we hypothesized a potential physical interaction between these components. To test this, we conducted yeast two-hybrid (Y2H) assays with N and C-terminus of CNGC13 with N and C-terminus of PEPR1 and PEPR2. No interaction of CNGC13 with PEPR1 was observed. However, we show an interaction of the cytosolic PEPR2 C-terminus (PEPR2-CT) with the cytosolic N-terminal of CNGC19 (CNGC19-NT) that was visible in SD-Leu/-Trp/-His (TDO) plate (**Figure 12A**). The kinase domain of PEPR2 is located at the C-terminus. To further validate this interaction, we employed bimolecular fluorescence complementation (BiFC) and co-expressed full length CNGC13-cYFP and PEPR2-nYFP, which yielded reconstituted YFP fluorescence at the plasma membrane, confirming the physical proximity of the two proteins (**Figure 12B**). We also investigated *in planta* interaction of PEPR2 with CNGC13 using split-luciferase complementation imaging assay and found similar observations (**Figure 12C**). Finally, we confirmed this with co-immunoprecipitation (Co-IP) assays in *Nicotiana benthamiana* following transient co-expression of CNGC13-FLAG and PEPR2-HA. CNGC13 co-immunoprecipitated PEPR2, suggesting a direct interaction at the plasma membrane (**Figure 12D**). These results prove that pattern recognition receptor, PEPR2 directly interacts with CNGC13 channel.

**Figure 12:**
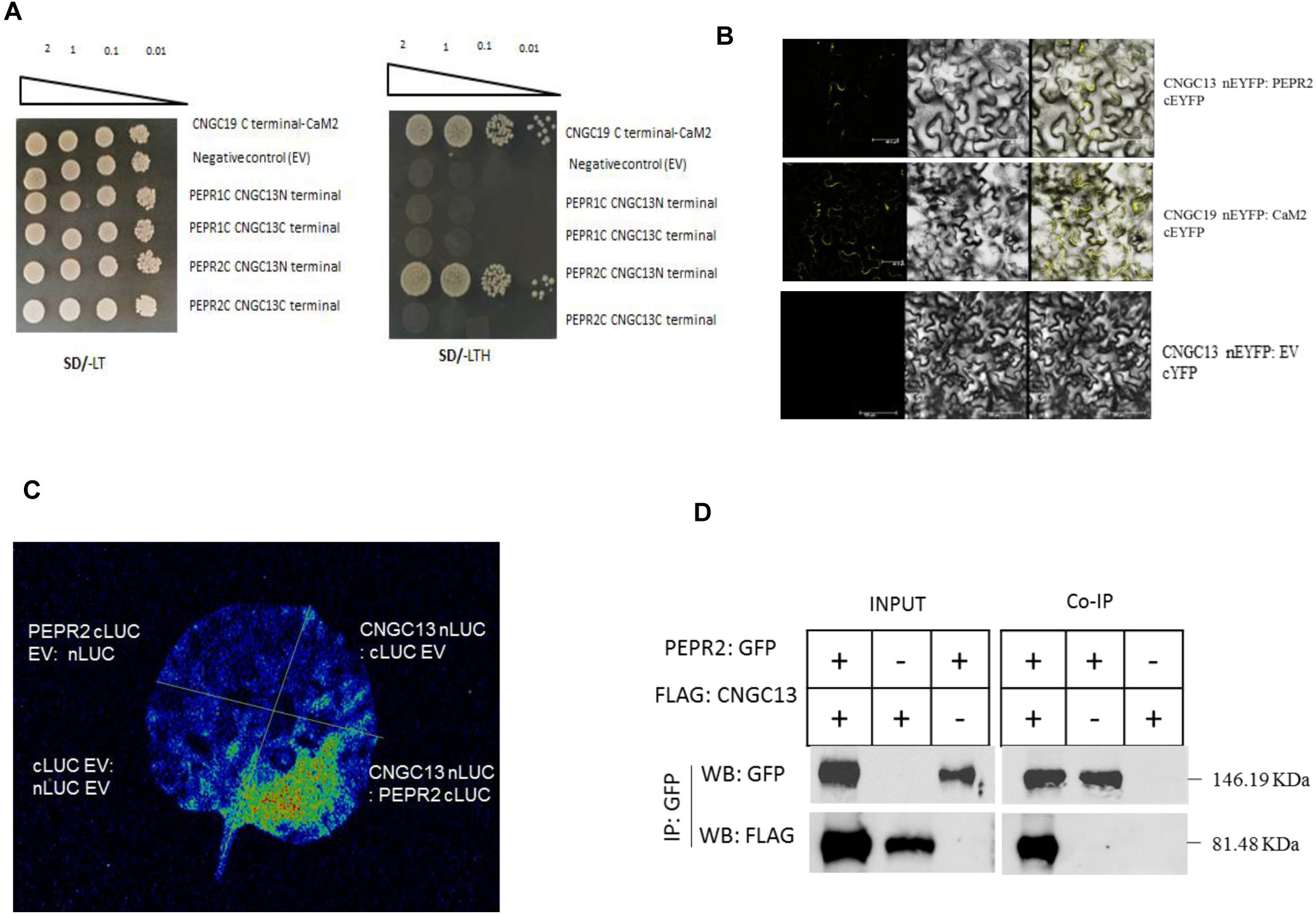
CNGC13 interacts with pattern recognition receptor, PEPR2. **(A**) Y2H interaction study of CNGC13 and PEPRs. Y2H gold yeast cells co-transformed with pGBKT7-CNGC13 N and C-terminal and pGADT7-PEPR1 and PEPR2 C-terminal were grown in non-selective SD-*Leu*/-*Trp* (DDO) medium to OD_600_ = 1. Ten μL from a 10-time dilution series were spotted on control non-selective DDO and selective TDO + 0.3mM 3-AT medium. (**B**) BIFC assay with full length CNGC13 and PEPR2, CNGC19 and CaM2 is used as positive control. CNGC13 fused with the N-terminal part of YFP and PEPR2 fused with the C-terminal part of YFP and transiently expressed in *N. benthamiana* by *Agrobacterium* mediated infiltration. Images shows the plasma membrane, and the superimposed images are of bright field and YFP. (**C**) Split luciferase assay was performed using full length CNGC13 fused with the pCAMBIA1300 nLUC and PEPR2 fused with the pCAMBIA1300 cLUC vector and transiently expressed in *N. benthamiana* by *Agrobacterium-*mediated infiltration. Images show only CNGC13 nLUC/ PEPR2 cLUC zone has stronger fluorescence than the other three control parts: nLUC/cLUC; CNGC13 nLUC/ cLUC; and nLUC/ PEPR2 cLUC. (**D**) Co-immunoprecipitation (Co-IP) assays showing interactions of CNGC13 with PEPR2. *proCNGC13:FLAG-CNGC13* and *35S:PEPR2:GFP* constructs were transiently expressed in *N. benthamiana* leaves. Protein extracts (input) were immunoprecipitated with α-GFP or α-FLAG antibody and resolved by SDS-PAGE. Protein-protein interactions were immunodetected using α-GFP and α-FLAG antibodies.

## Discussion

Insect herbivory rapidly triggers systemic signalling to activate defense in distal tissues. In Arabidopsis, early wound-induced signals include electrical, chemical, and hydraulic cues (Hedrich et al., 2023). Two parallel pathways underlie this response: (1) glutamate leakage and apoplastic pH changes activate Ca^2+^ channel, GLRs, triggering electrical signals and Ca^2+^ waves (Ngyuen et al., 2018; Toyota et al., 2018; Shao et al., 2020; Grenzi et al., 2023); thioglucosidases (TGGs) from phloem cells, catalyze glucosinolate breakdown into reactive aglucones that activate slow wave potentials (SWPs) and systemic Ca^2+^ waves (Gao et al., 2023). (2) hydraulic changes activate the mechanosensitive channel MSL10, leading to membrane depolarization and subsequent Ca²⁺ influx via H⁺-ATPase-mediated repolarization (Moe-Lange et al., 2021; Kumari et al., 2019). So far, only a few Ca^2+^-permeable channels are known to regulate systemic Ca^2+^ waves and also directly contribute to herbivore resistance and GLR3s (GLR3.1, GLR3.3, GLR3.6), the plasma membrane-localized CNGC19, the vacuolar two-pore channel TPC1, and Annexin1 are the only known members (Mousavi et al., 2013; Meena et al., 2019; Vincent et al., 2017; Ngyuen et al., 2018; Malarba et al., 2021; Kloth and Dicke, 2022). *At*GLR3.3 localize to the endoplasmic reticulum and plasma membrane, *At*GLR3.1 to endoplasmic reticulum, while AtGLR3.6 localizes to the tonoplast (Nguyen et al., 2018; Bellandi et al., 2022; Grenzi et al., 2023). Given the limited known Ca^2+^ channels and the intracellular localization of key GLRs, additional plasma membrane-localized channels likely contribute to systemic wound signalling. Plant cyclic nucleotide-gated channels (CNGCs) are tetrameric cation channels on the plasma membrane with a cytoplasmic N-terminal (NT) and C-terminal (CT) region per subunit, pore region (P loop) between S5 and S6 that permits ion transport (Jarratt-Barnham et al., 2021). Cryo-EM structures of Arabidopsis CNGC1 and CNGC5 revealed a unique extracellular domain stabilized by disulfide bonds, essential for channel gating through coupling of the voltage-sensing and pore domains and independent of cAMP/cGMP gating. The pore domain contains a selectivity filter with a constriction-site glutamine residue that confers Ca²⁺ selectivity (Wang et al., 2025). CNGCs form heterotetrameric complexes which may have unique functional characteristics, compared to homotetrameric channels (Dietrich et al., 2020).

We identify CNGC13 as a novel plasma membrane-localized Ca^2+^ channel essential for systemic wound signalling and anti-herbivore defense in *Arabidopsis thaliana*. CNGC13 is rapidly induced in both local and systemic leaves and is specifically expressed in phloem tissues, aligning with its role in vascular signal transmission. In wounded plants, phloem sieve elements and xylem contact cells coordinate long-distance electrical and Ca^2+^ signalling (Nguyen et al., 2018), and CNGC13 expression in phloem overlaps with GLR3.3, CNGC19, and MSL10, suggesting potential coordination among these channels. Electrophysiological analysis reveals that CNGC13 is activated by membrane hyperpolarization, similar to CNGC19, implicating it in the generation or propagation of the system potential, a self-propagating electrical signal initiated by wounding and herbivory (Zimmermann et al., 2009; 2016). While the upstream link between system potential and SWP, as well as downstream jasmonate induction, remains unclear, CNGC13 may bridge these processes. Although CNGC13 and CNGC19 are co-expressed in phloem and contribute synergistically to herbivore resistance, they do not physically interact or form heteromers. Instead, they exhibit transcriptional compensation and additive defense phenotypes in double mutants. Given the established role of CNGC19 in local Ca^2+^ influx and the upregulation of CNGC19 in *cngc13*, we propose that these channels form a distributed, context-dependent network of CNGC-mediated Ca^2+^ relays coordinating plant systemic immunity.

Loss of CNGC13 function impairs the velocity and propagation of systemic Ca^2+^ waves, particularly in distal leaves where signal transmission is confined to vascular tissues. In *cngc13* mutants, both TGG1 expression and isothiocyanate accumulation are reduced in systemic tissues, linking CNGC13 activity to wound-induced glucosinolate aglucone production. CNGC13 is the first Ca^2+^ channel known to functionally depend on the TGG-based signalling mechanism in response to mechanical wounding. Additionally, the initial hyperpolarization observed prior to Ca²⁺ elevation in systemic leaf of wild-type and *msl10-1* plants (Moe-Lange et al., 2021) suggests that CNGC13 which is activated by membrane hyperpolarization, is an important component of systemic signalling. Our data positions CNGC13 as a functionally essential component for TGG dependent vascular-dependent systemic signalling. The corresponding defect in systemic, but not local, jasmonate accumulation further supports its role in orchestrating distal hormone signalling. This systemic-specific phenotype is reminiscent of the defects observed in *glr3.3 glr3.6* and *msl10* mutants, suggesting that CNGC13 acts within a shared network of ion channels that operate in a vasculature-dependent manner to facilitate long-range jasmonate mediated immunity. CNGC13 and GLRs thus might mediate parallel branches of systemic wound signalling in *Arabidopsis*. The absence of coupling of SWP and Ca^2+^ elevation to long-distance jasmonate induction in non-vascular plants (Sanmartín et al., 2024), further highlights the specialized adaptation of vascular plants to herbivory.

Wounding leads to the release of various DAMPs, including peptides like *At*Peps, which are perceived by the receptor kinases PEPR1 and PEPR2 to activate immune responses (Krol et al., 2010; Tang et al., 2015; Prajapati et al., 2025). PEPR-mediated signalling has a critical role in amplifying PTI and coupling local and systemic immunity via jasmonate pathway in plant-microbe interactions (Ross et al., 2014). The vasculature localized receptors, PEPR1 and PEPR2 are strongly activated upon herbivore attack and *pepr1 pepr2* double mutant plants, display reduced resistance to *Spodoptera littoralis* larvae, reduced jasmonic acid levels and impaired external JA-Ile induced Ca^2+^ elevation (Klauser et al., 2015; Mittal et al., 2024). The JA responsive transcription factor, MYC2 also binds to the G-Box motif of *PEPR1* and *PEPR2* promoter and activates their expression (Mittal et al., 2024). No direct role of PEPR in wound mediated leaf-to-leaf systemic signalling is known so far. We demonstrate that *cngc13* mutants exhibit reduced cytosolic Ca^2+^ transients and ROS generation upon *At*Pep1 stimulation. CNGC13 responsiveness to *At*Peps may reflect a broader functional role in DAMP-induced Ca^2+^ influx, in a tissue-specific and herbivory-relevant context. The only other known channel in *At*Pep1 induced Ca^2+^ influx in Arabidopsis is CNGC2 via cGMP production (Qi et al., 2010). We show an interaction of PEPR2 C-terminus domain with N-terminus of CNGC13. The close spatial proximity of CNGCs and receptor kinases (RKs) at the plasma membrane suggests that RKs might play a central role in modulating CNGC activity through direct phosphorylation or scaffolded interactions. CNGC20 is the only other Ca²⁺ channel shown to directly interact with the receptor kinase BTL2, which phosphorylates the channel and activates it, when BAK1 is compromised (Yu et al., 2023). We previously reported a weak interaction between CNGC19 and the *e*ATP receptor DORN1 kinase domain (Kundu et al., 2025). Together, these findings support the existence of conserved receptor kinase–CNGC regulatory modules in plant immunity.

In summary, we identify CNGC13 as a key component of systemic wound signalling in *Arabidopsis*, functioning in the vasculature to coordinate TGG-dependent glucosinolate breakdown, Ca^2+^ wave propagation, jasmonate biosynthesis and herbivory resistance (**Figure 13**). CNGC13 and GLRs appear to define distinct but complementary branches of the signalling network where GLRs initiate rapid apoplastic responses, while CNGC13, acting downstream of PEPR2, amplifies Ca^2+^ signals to sustain the system potential that is required for long-distance signal propagation and defense activation. The physical interaction between PEPR2 and CNGC13 suggests a conserved mechanism by which receptor kinases regulate Ca^2+^ channels to link DAMP perception with long-distance defense activation. Future work will focus on the biochemical basis of vasculature localized PEPR2– CNGC13 interaction, effect on channel activity and structural determinants that enable receptor-mediated gating of CNGC13 and its role in systemic immunity.

**Figure 13:**
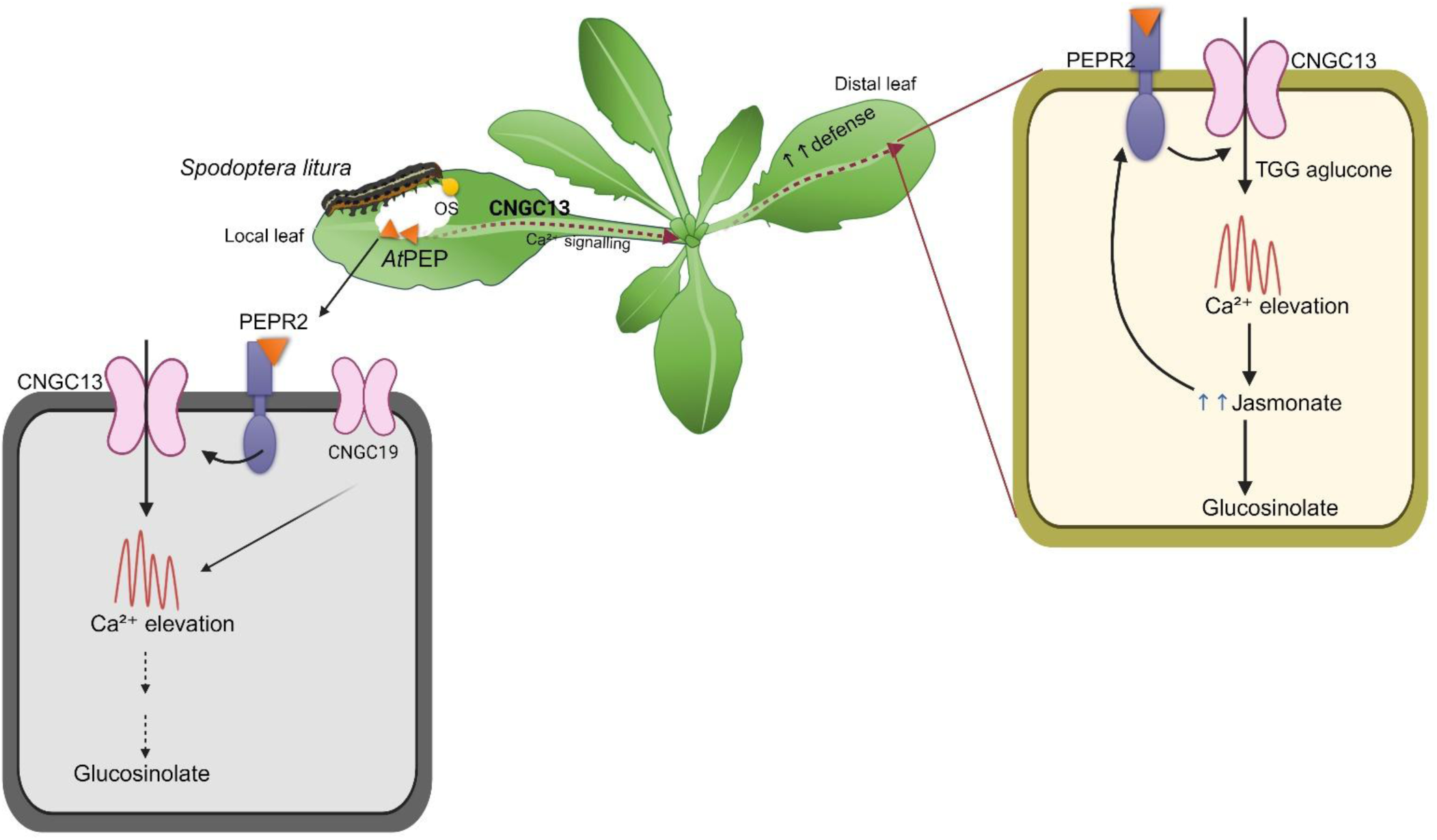
CNGC13 mediated modulation of plant immunity during insect herbivory. *Spodoptera litura* feeding on local leaves induces the release of endogenous elicitors such as *At*Peps, which are perceived by the receptor kinase PEPR2 at the plasma membrane. *S. litura* oral secretions (OS) also releases additional HAMPs and plant hormones that are sensed by unknown receptors in plants. *At*Pep receptor, PEPR2 activates CNGC13, a phloem-localized, plasma membrane Ca²[ channel, leading to cytosolic Ca²[ influx and downstream activation immune responses along with other HAMPs which culminates in glucosinolate biosynthesis in local leaf. CNGC13 and CNGC19 function cooperatively in herbivore defense, but do not physically interact, suggesting independent but converging roles on glucosinolate based plant defense in local leaf. In the systemic leaves, CNGC13 facilitates thioglucosidase (TGG)-dependent breakdown of aliphatic glucosinolates into reactive aglucones, which amplify Ca²[ signaling and promote jasmonate biosynthesis. Elevated jasmonates reinforce systemic defense against herbivory. CNGC13 operates in a distinct but converging pathway to GLRs to sustain long-distance Ca²[ wave propagation and immune activation.

## Materials and Methods

### Plant Material

*Arabidopsis thaliana* (ecotype: Columbia; Col-0) and T-DNA insertion mutant lines of *cngc13-1* (Salk_057742), *cngc13-2* (Salk_082668), and *cngc19-2* (Salk_129200) seeds were obtained from TAIR (Alonso *et al*., 2003). T-DNA insertion was confirmed by genotyping with LP/RP and Lb1.3/RP primer combination. CRISPR knockout mutant lines of CNGC13 (*cngc13-3*) were generated using the pKIR1.1 vector (Tsutsui and Higashiyama, 2017). Four sgRNAs targeting CNGC13 with minimal off-targets were designed using CRISPR GE, synthesized with ATTG overhangs, phosphorylated with T4 PNK, annealed, and ligated into *AaRI*-digested pKIR1.1 using T4 DNA ligase. Positive clones were used for the generation of transgenic lines via Agrobacterium-mediated transformation. Cas9-positive T1 seeds were selected by red fluorescence; Cas9-free lines in the T2 generation were identified by absence of fluorescence. Loss of function of *CNGC13* in the lines were confirmed by RNA-based genotyping (**Supplementary** Fig.1). A double mutant was generated by crossing *cngc13-1*\**cngc19-2* and the homozygous T3 generation plant was used for all experiments. Homozygous mutant seeds were checked for absence of *CNGC13* and *CNGC19* transcript using gene specific primers. *Arabidopsis* seeds were sown in pots containing a moist mixture of soilrite and agropeat (1:1) and kept for 2 days at 4°C under dark conditions for stratification. Stratified seeds were then transferred to ventilated growth chambers (Percival) with a short-day cycle (10-h-light/14-h-dark) with 60% humidity at 22°C under 150 µmol m^-2^s^-2^ light intensity. 4-5-week-old rosette plants were used for most of the experiments unless otherwise mentioned.

### Insect growth and Plant treatments

After *S. litura* neonates hatched from eggs, they were allowed to feed on agar based artificial diet (Bergomaz *et al*., 1986) under 23-25°C. For insect biomass assay individual 2^nd^ instar larvae having equal weight were selected. Three larvae were placed on each plant and covered with perforated plastic bags. After 8 days of feeding, larvae were weighed again for calculating insect weight gain. Each experiment was performed with 12 plants and the experiment was repeated for 3 times. For GS estimation or phytohormone quantification experiment, feeding was performed with 12 h pre-starved 4^th^ instar larvae. For experiments with insect oral secretions (OS), 4^th^ instar *S. litura* larvae reared on artificial diet were fed on Arabidopsis leaves for 24 h prior to collecting OS on ice. The OS was centrifuged at 13,000 rpm for 2 min and diluted 1:1 with water. For simulated herbivory experiment, wounding was done with a pattern wheel (6 vertical motions) on either side of the leaf and total of 20 µL of fresh diluted OS or water (control) was spread across all the holes on a single leaf. For expression analysis in systemic leaves, leaves were numbered, the petiole of leaf number 8 was cut with scissors. Systemic leaf 13 with vascular connection to the leaf number 8 was harvested after different time points for expression analysis (Kiep *et al.,* 2015; Mousavi *et al.,* 2013).

### Real-Time Quantitative PCR

Plant samples were harvested, immediately frozen in liquid nitrogen and stored at −80°C until further processing. Tissue samples were ground in liquid nitrogen and total RNA was isolated using Trizol method (Invitrogen) according to the manufacturer’s protocol. Total RNA was treated with DNaseI to remove any DNA contamination. RNA quantity and quality was analysed with Nanodrop (Thermo Scientific). 1µg of total RNA was used to synthesize cDNA with a mix of oligo-dT_18_ primers using the Omniscript cDNA synthesis kit using the manufacturer’s protocol (Invitrogen). Primers from exon-exon junction were designed by NCBI primer design tool (http://www.ncbi.nlm.nih.gov/tools/primer-blast) with 100-150 bp amplicon size. RT-qPCR was done in 96 well semi-skirted plates (Biorad) with 50 ng cDNA in iTaq Universal SYBR green supermix (Biorad). *RPS18* was used as endogenous control to normalize transcript level. Fold change of target genes were calculated with the ΔΔCT equation (Rao *et al*., 2013) and relative to the mRNA level of target genes in the control leaf, which were defined as 1.0. All of the experiments were run with at least four biological replicates. The primer pairs used are listed in **Supplemental Table 1**.

### Vector construction and expression of YFP/GFP Fusion Protein

The coding sequence of *CNGC13* was cloned into the pEG101 to generate a C-terminal YFP fusion under the control of a Cauliflower Mosaic Virus promoter (CaMV35S) using gateway cloning technology (Invitrogen). The attb-flanked PCR products were recombined into the donor vector pDONR207 using BP clonase. Sequence-verified entry clones were recombined with pEG101 using LR clonase reaction. The resulting *CNGC13-YFP* construct was transformed into the *Agrobacterium tumefaciens* strain GV3101. For transient expression in *Nicotiana benthamiana*, overnight bacterial cultures were resuspended in Infiltration buffer (10 mM MES-KOH, pH 5.6; 10 mM MgCl2; 150 uM acetosyringone) to an OD_600_ of 0.8. Equal volumes of *Agrobacterium* carrying *CNGC13-YFP*, a plasma membrane marker (PM-mCherry) and the p19 silencing suppressor were mixed (1:1:1), incubated at room temperature for 3 h, and used for leaf infiltration 2 days later, infiltrated leaves were imaged using a TCS SP5 confocal laser scanning microscope (Leica Microsystems) with excitation/emission wavelengths of 514-527 nm (YFP) and 587-610 nm (mCherry).

Subcellular fractionation was performed using the Qproteome Cell Compartment Kit (Qiagen) according to the manufacturer’s instructions. Protein fractions were analysed by immunoblotting with anti-GFP primary antibody (Agrisera Antibodies, Sweden) and then with goat-anti rabbit IgG-HRP conjugates (Merck) followed by chemiluminescence with clarity TM western ECL substrate (Bio-Rad). For stable transformation, the *proCNGC13:gCNGC13:GFP* construct (synthesized by GeneArt) in the pCAMBIA1300 vector was introduced into the *Arabidopsis thaliana* Col-0 plants using the floral dip method and homozygous T3 generation plants used for experiments. The TCS SP8 confocal laser scanning microscope (Leica Microsystem) was used to visualise CNGC13-GFP in roots stained with propidium iodide (PI) for 5 min. PI excitation and emission were visualised at wavelengths of 561 and 615 nm respectively. For sieve plate imaging the roots were stained with aniline blue dye (0.1% [w/v]) for 30 mins and visualised by epifluorecence microscope using a short-wavelength emission filter set (330 to 385-nm excitation; 420-nm emission) or enhanced GFP (470 to 490-nm excitation; 515-nm emission).

### *CNGC13* promoter assay

For promoter analysis of *CNGC13*, a 2.5-kb promoter region upstream of the start codon was amplified with Phusion Taq polymerase (Thermo-Fisher) using *Arabidopsis* genomic DNA. Amplified promoter product was cloned in pCAMBIA1304 vector using *XbaI*/*NcoI* restriction sites. For GUS reporter assay, 12 days old transgenic seedlings of *ProCNGC13:GUS* were either locally wounded, systemically wounded or treated with 100 nM *At*Pep1 or *At*Pep2 or water (control) for 2 h. After 2 h seedlings were harvested and placed in GUS solution (50 mM of sodium phosphate buffer at pH 7.0, 2 mM of EDTA (w/v), 0.12% Triton, 0.4 mM of iron ferrocyanide, 0.4 mM of iron ferricyanide, and 1.0 mM of 5-bromo-4-chloro-3-indolyl-β-D-glucuronide cyclohexylammonium salt) and incubated for 4 h in the dark at 37°C. Chlorophyll was removed by treating the seedlings with 70% ethanol at 65^○^C. Seedlings were photographed under bright field microscope.

### Vector construction of electrophysiology

For electrophysiology experiment full length *CNGC13-YFP* CDS was cloned into *Xenopus laevis* (the African clawed frog) oocyte expression vector. Since, *CNGC13* is a plant gene and we are expressing it in a completely different animal system, for better expression *CNGC13-YFP* cassette was codon optimized for *Xenopus* oocytes using codon optimizer tool available on Thermo-Scientific system (https://www.thermofisher.com/in/en/home/life-science/cloning/gene-synthesis/geneart-gene-synthesis/geneoptimizer.html). For restriction-based cloning, *EcoRI* and *BamHI* restriction enzyme sites were added in codon optimized *CNGC13-YFP* sequence. Complete cassette of *CNGC13-YFP* sequence along with restriction enzyme sites was artificially synthesized in the pUC57 vector by GeneArt. The *CNGC13-YFP* (2844 bp) cassette was removed from pUC57 vector by using *EcoRI*, *BamHI* and *ScaI* enzymes. *CNGC13-YFP* (2844 bp) cassette was cloned into the pGEMHE-EV (3022 bp) vector. Positive clones were confirmed by restriction digestion and sequencing. The primer pairs used are listed in **Supplemental Table 1**.

### Confocal Microscopy to detect expression and localization of CNGC13-YFP in *Xenopus laevis* oocytes

Stage 5 and 6 oocytes from adult female *Xenopus laevis* (the African clawed frog) were surgically removed from anesthetized frogs in accordance with a protocol approved by the Institutional Animal Ethics Committee (IAEC), IISER Bhopal. Oocytes were digested with collagenase type 2 enzyme (1 mg/mL) in calcium-free OR2 buffer containing NaCl (82.5 mM), KCl (2.5 mM), MgCl_2_ (1 mM) and HEPES (5 mM) at pH 7.6 for 1 h at 18° C on an automated rocker and then de-folliculated with cut pipettes. Capped RNA was prepared from the linearized plasmid DNA templates using the HiScribe™ T7 High Yield RNA Synthesis Kit (New England Biolabs), following the manufacturer’s instructions. De-folliculated oocytes were injected with *CNGC13-YFP* cRNA solution (50 nL) and stored at 18° C in ND96 solution containing NaCl (96 mM), KCl (2 mM), HEPES (5 mM), MgCl_2_ (1 mM), CaCl_2_ (1.8 mM) and gentamycin (50 µg/mL) at pH 7.6. Un-injected and *CNGC13-YFP* cRNA injected oocytes (72 to 96 h post-injection) were imaged in bright field and fluorescence confocal mode. Oocytes were imaged with a 10.0X objective on a FV3000 Confocal Microscope (Olympus). YFP was excited using the 488 nm line at 1.4% transmissivity with a PMT voltage of 640 V while the emission was recorded from 500 to 550 nm. Line scans were performed on fluorescence images of both un-injected and CNGC13-YFP cRNA injected oocytes using ImageJ software.

### Activity assays of CNGC13 channels by two-electrode voltage clamp recordings

TEVC recordings for CNGC13-YFP RNA injected oocytes were performed 72 to 96 h post-injection. The recorded data were filtered at 1 kHz and sampled at 5 kHz. Recording microelectrode resistances were between at 0.1-1 MΩ when filled with 3 M KCl solution. Two separate recording buffers were prepared—one for recordings in presence of Ca^2+^ ions and the other for those in the absence of Ca^2+^. The former contained CaCl_2_ (15 mM), mannitol (200 mM) in HEPES-Tris (10 mM) at pH 7.4 and the latter contained KCl (10 mM), mannitol (200 mM) in HEPES-Tris (10 mM) at pH 7.4 (Meena, Prajapati et al. 2019). Stock solution of dibutyryl c-AMP was made in mannitol (200 mM) in HEPES-Tris (10 mM) buffer at pH 7.4. Before and after compound administration, the oocytes were subjected to a step-protocol wherein they were held at −50 mV for 100 ms, and then the voltage was stepped up in +20 mV increments between −130 mV to +30 mV for 2 sec before stepping back to the holding voltage for 500 ms.

### Complementation of *cch1*/*mid1* yeast mutant with CNGC13

The yeast wild type and calcium channel mutant *cch1*/*mid1* strains were transformed with pYES2 empty vector and *pYES2-CNGC13* constructs by Clontech Yeast Transformation Kit (Clontech Laboratories Inc). Transformants were selected on yeast minimal medium lacking uracil. The complementation assay was performed according to previously reported methods (Ali *et al*., 2006; Charpentier *et al*., 2016). Cells were grown in selective liquid medium lacking uracil over-night, washed 3 times with sterilized Milli Q water and resuspended in liquid YPD medium to an OD_600nm_ of 4. YPD medium contains galactose as a carbon source to induce expression of CNGC13 by *GAL1* promoter. An aliquot (100 µl) of this resuspension was added to 4 ml of top agar (YPD with 0.7% agar), mixed and overlaid on YPD agar plates. Immediately after solidification of top agar-containing cells, sterile cellulose discs (6mm diameter and 45 µm pore size) containing 20 µg of synthetic alpha factor (Sigma T6901) were placed on the nascent lawn. Plates were incubated at 30° C for 2 days

### Ca^2+^ measurement using Aequorin reporters

A stable transgenic Arabidopsis line expressing cytosolic aequorin in the *cngc13-1* background was developed using the floral dip method using pMAQ2 construct. Stable transgenic of T3 generation of homozygous plants with good discharge were used for Ca^2+^ measurement. CRISPR lines of CNGC13 (*cngc13-3*) was generated in aequorin background as explained earlier with WT. *Arabidopsis* seedlings were grown in liquid in 24 well plate containing half strength MS media under short day condition (14 h/10 h). Seedlings were incubated in dark for overnight in 96 well luminometer plates (Thermo Fisher) containing 5 µM coelenterazine (PJK international) at 21°C. Bioluminescence counts were recorded as relative light unit using luminometer (Luminoskan Ascent, v2.6; Thermo Fisher Scientific). After background reading of 120 s, 100 nM of *At*Pep1 or *At*Pep2 (GenScript), or water (control) was added manually and signals were measured as relative light units per s for 300 s with 1 s integration and 1 s interval time. At the end, signals were normalized by discharging remaining aequorin of the seedling with 2M CaCl_2_ in 20% ethanol. Bioluminescence counts were converted into cytosolic Ca^2+^ concentration according to Rentel and Knight (2004).

### Whole plant real-time Ca^2+^ imaging using GCaMP3 reporters

Transgenic *Arabidopsis thaliana* lines expressing GCaMP3 in wild-type (WT_GCaMP3_) and *cngc13-1 and cngc13-3* (*cngc13*_GCaMP3_) backgrounds were used for live Ca^2+^ imaging. Plants were grown on soil for 2 weeks under the short day conditions (8 h/16 h) prior to experimentation. The *cngc13*_GCaMP3_ line was generated by floral dipping using a modified pCAMBIA2300 binary vector containing *2X CaMV35S::GCaMP3*. Experiment was performed in the T3 generation. To ensure comparable GCaMP3 dynamic range among genotypes, both WT_GCaMP3_ and *cngc13*_GCaMP3_ leaf discs were treated with 2M CaCl_2_ and 20% ethanol following each experiment. WT_GCaMP3_ and *cngc13*_GCaMP3_ displaying comparable GCaMP3 dynamic ranges were used, thereby minimising difference in the fluorescence signal that could affect Ca^2+^ response comparisons. For all the experiments 12-15-day-old plant was used, cutting leaf 1 petiole caused a systemic Ca^2+^ increase that propagated within the rosette of intact soil-grown plants. The fluorescence images were captured every 5 s for a total of 5 min. Arabidopsis GCaMP3 plants were imaged with a Nikon SMZ18 stereomicroscope (http://www.nikon.com/) equipped with an ORCA-Fusion BT sCMOS camera (https://www.hamamatsu.com/eu/en/product/cameras/cmos-cameras/C11440-42U40.html). Excitation light (460 nm) was produced by a CoolLED pE-300ultra (https://www.coolled.com/products/pe-300ultra/) with GFP filter cube to collect GCaMP3 emission. A Plan Apo 0.5X objective was used, and images were collected with an exposure time of 0.5 s with a 2 x 2 camera binning (1440 x 1024 pixels). GCaMP3 images were acquired every 5 sec with the GFP filter cube. Mean fluorescence of images was quantified by using ImageJ/Fiji (National Institute of Health, USA). TIFF files were used for data processing and whole leaf mean fluorescence excluding cut region was calculated by selecting leaf area using freehand selection tool (**Supplementary Figure 3B**). Normalized fluorescence intensity (ΔF/F_0_) was calculated according to ΔF/F_0_ = F-F_0_/F_0_ (F-fluorescence after wounding, F_0_ - basal fluorescence intensity). For the analysis of Ca^2+^ wave velocity, a significant rise in signal within each Region Of Interest (ROI) was defined as the point at which the fluorescence intensity exceeds two times the standard deviation (2X SD) above the pre-stimulation baseline (F_0_). The 2X SD threshold was calculated from the F_0_ data using Image J software. The time points at which the Ca^2+^ signal increased in ROI1 to ROI2, ROI3 and ROI4 were identified, and the time-lag (t = t2-t1) between the two ROIs was calculated. The physical distance between ROI1 and ROI2 was subsequently measured, and the velocity of the Ca^2+^ wave was determined by dividing this distance by the time lag (Uemura et al., 2021).

### ROS measurement

Leaf disc of *Arabidopsis* rosette of WT and *cngc13* mutants were prepared from different plants. Single leaf disc per well was transferred to 100 µl of milli-Q water in a black 96 well plate well and incubated in plant growth chamber at 22^°^C and 60% relative humidity in dark. Horseradish peroxidase (Sigma-Aldrich) was dissolved in water to make stock of 10 mg/ml. Luminol (Sigma-Aldrich) was also dissolved in water to make final concentration of 0.5 mM. Imaging solution was prepared by diluting HRP (20 µg/ml) luminol (1 µM) and 100 nM *At*Pep in water. Next day plants were placed in luminometer (Luminoskan Ascent, v2.6; Thermo Fisher Scientific) and allowed to acclimatize for 10 min. Water was replaced with 100 µl imaging solution and reading were taken in 8-well kinetic mode with 1 sec integration time and 1 sec interval time. Experiments were done with 3 independent biological replicates as per figure legends

### Glucosinolate Quantification

5-week-old whole plant rosette was immediately harvested in liquid nitrogen and freeze dried in lyophilizer. After grinding to fine powder, twenty-five milligrams of freeze-dried sample per plant was used for GS extraction. GS extraction was done in 1 mL of 80% methanol solvent containing 50 µM intact 4-hydroxybenzylglucosinolate (sinalbin) as internal standard. After centrifugation, 800 µL supernatant was loaded onto pre-conditioned DEAE-Sephadex A25 columns. Column was washed with washing solution and treated with 30µl sulfatase (Sigma-Aldrich) for de-sulfation. After overnight incubation GS were eluted with 0.5 ml HPLC grade water. Eluted desulfoglucosinolates were separated using HPLC (Agilent 1100 HPLC system; Agilent Technologies) on a reverse-phase C-18 column (Chromolith Performance RP18e, 100 × 4.6 mm; Merck) with a water-acetonitrile gradient. Experiments were done with 3 independent biological replicates as per figure legends

### Glucosinolate and Isothiocyanate quantification with extracted midrib

Five-week-old *Arabidopsis* leaves were numbered, with leaf 8 selected for wounding and the midrib of leaf 13 collected for metabolite analysis. Wounding was performed on leaf 8 using ridged forceps, pressed four times in parallel to create uniform damage covering approximately 40% of the leaf area, following Gao et al. (2023). Equal pressure was applied, and each leaf was wounded within 10 seconds. The midrib of leaf 13 was collected after 2 min of wounding, pooled from 4–6 plants per replicate, and immediately flash-frozen using liquid nitrogen. Frozen tissues were pre-weighed and ground in liquid nitrogen, and metabolites were extracted using 500 µL of 70% methanol containing 0.1% formic acid and 50 µM Sinalbin (Sigma) as an internal standard. Samples were vortexed for 2 min and incubated on ice for 2 min; this cycle was repeated five times. Extracts were centrifuged at 12,000 rpm for 20 minutes at 4°C. Glucosinolate analysis was performed as described in Gao et al. (2023) For detection, the mass spectrometer was operated in positive ion mode for intact glucosinolates and in negative ion mode for isothiocyanates. Integrated peak areas were normalized to both the initial plant mass and the sinalbin internal standard (usually used for GS analysis as std), deviating from the original protocol in Gao et al., 2023. Experiments were performed using three independent biological replicates, as indicated in the figure legends.

### Phytohormone quantification

Phytohormone levels were measured from systemic leaf 13, collected 15 min after cutting the petiole of leaf 8 from 4-week-old *Arabidopsis* plants. Each replicate is a pool of 3 leaves collected from 3 different plants (200–250 mg). For local responses, two rosette leaves (200–250 mg) from wounded or herbivore fed 4-week-old plants were harvested, flash-frozen in liquid nitrogen, and stored at – 80°C until processing. Tissue samples were finely ground in liquid nitrogen using mortar pastel and phytohormone was extracted with 1.5 mL of methanol containing 40 ng d_6_-jasmonic acid (HPC Standards GmbH, Cunnersdorf, Germany), 60 ng salicylic acid-d_4_ (Santa Cruz Biotechnology), 60 ng abscisic acid-d_6_ (Toronto Research Chemicals, Toronto, ON. CA) and 12 ng d_6_-jasmonic acid-isoleucine conjugate (HPC Standards GmbH) as internal standards. Samples were vortexed briefly for about 2 min followed by end to end shaking on horizontal tube rotator at 4°C for 20 min. Samples were than centrifuged at 12,000 x g at 4oC for 20 minutes. Supernatant was collected in a new tube and dried in a speed-vac system at room temperature. When samples were completely dried 500 µl fresh methanol (without internal standard) was added and vortexed well. Samples were then centrifuged at 16,000 x g for 5 min at 4°C. Samples were filtered using 0.45 µM filter (Merck) and quantified on a QTRAP6500 LC-MS/MS system (AB Sciex) as described (Vadassery *et al*., 2012). The absolute quantification of hormones were done using deuterated internal standards. Since it was observed that both the D6-labeled JA and D6-labeled JA-Ile contained 40% of the corresponding D5-labeled compounds, the sum of the peak areas of D5- and D6-compound was used for quantification. Experiments were done with 3 independent biological replicates as per figure legends.

### Yeast two hybrid (Y2H)

*PEPR1* and *PEPR2* C-terminal and *CNGC19* N and C-terminal were cloned into pGADT7-AD and *CNGC13-N* and *CNGC13-C* terminal were cloned into pGBKT7-BD respectively (Clontech Laboratories Inc). All constructs were made using the restriction based cloning. The pGBKT7-BD and PGADT7-AD constructs were co-transformed into Y2H Gold strain of yeast using Matchmaker Gold yeast Two-Hybrid system kit (Clontech Laboratories Inc), and selected for transformed cells on DDO (-LT) plates lacking leucine and tryptophan and were confirmed by colony PCR. Y2H Gold strain co-transformed with the empty vectors, pGBKT7-BD and pGADT7-AD were also used as a negative control. Transformed cells were grown in liquid double dropout media (-LT) for 2 days, adjusted to an OD_600_ of 2, 1, 0.1, 0.01, and 10 µl of each dilution were spotted on selection plates; DDO (-LT) and without histidine TDO (-LTH) + 0.3mM 3-AT (3-Amino-1,2,4-Triazol). Plates were grown at 30°C for 3 days. The primer pairs used are listed in **Supplemental Table 1**.

### BiFC Assay

*35s::nEYFP:CNGC13* and *35s::PEPR2:cEYFP* constructs were generated using an LR (Invitrogen) reaction between pDONR vector carrying both these inserts into destination vectors *pSITE-nEYFP-C1* (CD3 1648) and *pSITE-cEYFP-N1* (CD3 1651) respectively. The successful constructs were retransformed into *Agrobacterium GV3101.* Transformant cultures grown at (OD_600_) 1 and incubated in infiltration buffer (10 mM MES pH[5.6, 10 mM MgCl_2_, 100 μM acetosyringone) as described by Schütze *et al*. (2009) for 2.5 h at room temperature. The cultures were infiltrated into 5-week-old *Nicotiana benthamiana* leaves at a ratio of 1:1:1 along with p19, the silencing suppressor. The plants were then incubated at dark for 40-44 h. eYFP fluorescent signals were observed and captured from 44 to 48 h using confocal microscopy in Leica-SP8 systems. The primer pairs used are listed in **Supplemental Table 1**.

### Split-Luciferase Complementation Imaging Assay

*35S:CNGC13* and *35S:PEPR2* full length were cloned into pCAMBIA-nLUC and pCAMBIA-cLUC respectively. The primers have been listed in the **Supplementary Table S1**. The constructs were transformed into *Agrobacterium* (*GV3101* strain) and the culture was harvested by centrifugation at 4500 rpm for 15 min. The cultures were re-suspended in infiltration medium (1 M MgCl_2_, 100 mM acetosyringone, 1 M MES-KOH, pH 5.7) and incubated for 3 h in a dark at room temperature. The cultures were infiltrated in the abaxial surface of *Nicotiana benthamiana* leaves. Two days post inoculation the leaves were infiltrated with 1 mM luciferin and kept in the dark for 7 min. The infiltrated leaf samples were analyzed using low light cooled CCD imaging system (Bio-Rad). For Luciferase imaging, three independent experiments (with 6-8 plants per experiment) were conducted, and similar results were obtained.

### Co-immunoprecipitation assay

Transient expression of *proCNGC13:CNGC13:FLAG* and *35S:PEPR2:GFP* constructs was performed in *Nicotiana benthamiana* leaves for 48 h. Following expression, total proteins was extracted using an extraction buffer buffer (50 mM Tris-HCl pH 7.5, 100 mM NaCl, 10% glycerol, 5mM DTT, 1% protease inhibitor cocktail, 0.5% (v/v) IGEPAL, 1 mM Na_2_MoO_4_.2H_2_O, 1 mM NaF, 1.5 mM EDTA and 1.5M m Na_3_VO_4_). For each sample, 500 mg of infilterated leaf tissue was ground in the extraction buffer and incubated at 4°C for 90 min with gentle rotation for plasma membrane solubilisation. The homogenates were then centrifuged at 15,000 x g for 20 min at 4°C, and the resulting supernatant were collected. The extracted proteins were stored at −80°C until further use. Co-immunoprecipitation (Co-IP) was carried out using GFP-Trap® Magnetic Agarose Kit (Chromotek). Bead storage buffer was first removed, and the beads were equilibrated by washing with the extraction buffer. The washed beads were then incubated with the protein extracts for 6 h at 4°C with continuous rolling. Following incubation, the beads were washed 5-6 times with the extractiuon buffer to remove non-specifically bound proteins. Bound proteins were eluted using 2X SDS sample buffer. Eluted proteins were run on the SDS PAGE and the proteins were detected using α-GFP, and α-FLAG antibodies.

### Statistical Analysis

Statistical differences between different groups were detected by one-way ANOVA and a posthoc SNK test/Tukey Test or *t*-test in SigmaStat 2.03. Different letters indicate significant difference between treatments.

### Accession Numbers

AT4G01010 (CNGC13), AT1G17750 (PEPR2), AT3G17690 (CNGC19)

## Acknowledgements

We acknowledge the Department of Biotechnology (DBT), Government of India, NIPGR core grant, Janaki Ammal National Bioscience Award (HRD-17014/2/2023-HRD-DBT) and EMBO Global Investigator Network for funding the work in JV lab. We acknowledge student funding from: DBT-JRF for RP, UGC-NET / JRF (191620185954) for MK, DBT-SRF (3-504/2022-23) for MP. We acknowledge the Ministero dell’Istruzione, dell’Università e della Ricerca — Fondo per Progetti di Ricerca di Rilevante Interesse Nazionale 2022 (PRIN 2022NMSFHN) (to AC), and the Agritech National Research Center, funded by the European Union Next Generation EU (Piano Nazionale di Ripresa e Resilienza (PNRR) — Missione 4, Componente 2, Investimento 1.4 – D.D. 1032 17/06/2022, CN00000022) (to AC and BMOM). We thank Bhawana Sharma (model illustration), Kushboo Sharma (metabolomics), Shruti Mishra (aequorin), and former members Mukesh Kumar Meena and Riya Joon (seed and cloning resources). We acknowledge NIPGR Metabolome facility (funded by DBT (BT/INF/22/SP28268/2018) for phytohormone and metabolite quantification. We also acknowledge the NIPGR central instrumentation, phytotron facility and DBT-eLibrary Consortium (DeLCON) for providing access to e-resources, and the NOLIMITS Imaging Center of Excellence for Plant Biology and Other Life Sciences, established by the University of Milan. We thank the DST-FIST facility at IISER Bhopal for confocal microscopy support and Manas (Jeet Kalia lab) for help with Xenopus laevis oocyte imaging.

## Supplemental Data

**Supplemental Figure 1.** T-DNA insertion/CRISPR mediated mutation and genotyping of *cngc13* mutants

**Supplemental Figure 2.** CNGC13 is a plasma membrane localized Ca^2+^-permeable channel.

**Supplemental Figure 3.** GCaMP3 sensor dynamic range in WT and *cngc13* mutants

**Supplemental Figure 4.** CNGC13 is involved in generation of systemic Ca^2+^ elevation upon wounding

**Supplemental Figure 5.** Total Glucosinolate content in systemic veins after wounding

**Supplemental Figure 6.** Defense related phytohormone level in WT and *cngc13* mutants

**Supplemental Figure 7.** Jasmonate accumulation in local leaf after wounding is unaltered

**Supplemental Figure 8.** CNGC13 important for *At*Pep signalling

**Supplemental Figure 9.** Herbivory specific *CNGC* expression in mutants and CNGC 13 homodimer formation

**Supplemental Table 1. Primer List**

**Movie S1**: Systemic Ca^2+^ signals in WT GCaMP3 plant

**Movie S2**: Systemic Ca^2+^ signals in GCaMP3-*cngc13-1* mutant plant

**Movie S3**: Systemic Ca^2+^ signals in GCaMP3-*cngc13-3* mutant plant

