## Supplemental Table1 for "The Ca^2+^ Channel CYCLIC NUCLEOTIDE GATED CHANNEL13 (CNGC13) regulates systemic wound signalling and immunity against Spodoptera herbivory"

**Supplementary Table 1**

**Real time PCR primers**

| *CNGC13-RT-F* | CGCAACAATCGTGTAAGGTTCA |
| --- | --- |
| *CNGC13-RT-R* | AGTCCTCTACGAACATTCTTCAA |
| *RPS18-RT-F* | GCGTGTGCTCAACACTAACG |
| *RPS18-RT-R* | CTCACCAGCCCTCTTGTTCA |
| *TGG1 RT FP* | CGTTGATGTTTACAGGACGAAA |
| *TGG1 RT RP* | CAGTTGCATCTTTGCTCTCTTG |
| *TGG2 RT FP* | TGGCAGAAAGATCTAGACGTGA |
| *TGG2 RT FP* | TTCCTTTTGGAAGGATTCTTGA |
| *GLR3.3 RT FP* | AAGTGGCTGGGGATTTGCCTTTC |
| *GLR3.3 RT RP* | TTGATCTCTTGCAATGGCATCAT |
| *GLR3.6 RT FP* | GGGGATTTGCATTTCCGCGT |
| *GLR3.6 RT RP* | TCTATCTCTGCGCCTTGCAG |
| *JAZ10 RT FP* | ATCCCGATTTCTCCGGTCCA |
| *JAZ10 RT RP* | CTTTCTCCTTGCGATGGGAAGA |
| *LOX6 RT FP* | TGTGAACCACTGGCTAAGGAC |
| *LOX6 RT RP* | CTTTTACGCGCACGAGCATTG |
| *CNGC19 RT F* | CAAGTTACTCAGTGGGTGAGGT |
| *CNGC19 RT R* | GCATCATTTCCATCCTCCTTCG |

**Cloning and sequencing primers**

| *CNGC13-SeqF_1.2Kb* | TGTTTTCGGAAGAGGCGAGT |
| --- | --- |
| *CNGC13-SeqR_1.2Kb* | CATGTGAGAAGATCTTCGCC |
| *CNGC13-attB1-FP2* | GGGGACAAGTTTGTACAAAAAAGCAGGCTTCATGGCTTTTGGCCGC |
| *CNGC13-attB2-RP2* | GGGGACCACTTTGTACAAGAAAGCTGGGTTAGGGTTTCGGAGACTGAA |
| *CNGC13-attB2-C RP2* | GGGGACCACTTTGTACAAGAAAGCTGGGTTTTAAGGGTTTCGGAGACTGAA |
| *pCNGC13-2.5kb FP* | TCTAGAAGACGAAAAGGACGCA |
| *PEPR2_attB1-FP1* | GGGGACAAGTTTGTACAAAAAAGCAGGCTTCATGAGGAATCTTGGGTTACTCGAAATTAC |
| *PEPR2_attB2-RP1* | GGGGACCACTTTGTACAAGAAAGCTGGGTTCTAGTGAACTGAACCCGAAGTGCT |
| *PEPR2_attB2-C-RP2* | GGGGACCACTTTGTACAAGAAAGCTGGGTTGTGAACTGAACCCGAAGTGCT |
| *pCNGC13-2.5kb RP* | CCATGGGAGTCAAGCTTTTTTTG |
| *M13F* | CCCAGTCACGACGTTGTAAAACG |
| *M13R* | AGCGGATAACAATTTCACACAGG |
| *pCNGC13-2.5kb RP* | CCATGGGAGTCAAGCTTTTTTTG |
| *CNGC13/GUS FP* | CCATGGCTTTTGGCCGCAACA |
| *mGFP5 RP* | CAACAAGAATTGGGACAACTCC |
| *CaMV 35s FP* | GCA CCC CAG GCT TTA CAC TTT |
| *CNGC13 Seq FP* | TGTTTTCGGAAGAGGCGAGTTG |
| *CNGC13 Seq RP* | CTAATGCCCATGTGAGAAGATC |
| *CNGC13 Sq F* | AACTGATGAGCGCCACCACAAACGGT |
| *CNGC13 Sq R* | TTTAGGAGGTAGCAAAGGTAGGTTG |
| *CNGC13 YES2 FP* | CGGGATCCAACACAATGTCTGCTTTTGGCCGCAACAATC |
| *CNGC13 YES2 FP* | CGGGATCCAACACAATGTCTGCTTTTGGCCGCAACAATC |
| *CNGC19-C Y FP* | GGAATTCCATATGTCACTTGATCGAAGGAGGAT |
| *CNGC19-C Y RP* | CGGGATCCTTAACGGTTGGAATTGGAGTGAGCA |
| *CNGC19-N Y FP* | GGAATTCCATATGGCTCACACTAGGACT |
| *CNGC19-N Y RP* | CGGGATCCTTATTGAACAAATTTGGAATGAG |
| *CNGC13-C Y FP* | GGAATTCCATATGCAGAAATACTTGGAAT |
| *CNGC13-C Y RP* | GGAATTCTTAAGGGTTTCGGAGACTGA |
| *CNGC13-N Y FP* | GGAATTCCATATGGCTTTTGGCCGCAACA |
| *CNGC13-N Y RP* | GGAATTCTTAGTTCCAGTTCTGGAAAAAA |
| *CAM2Y-FP* | GGAATTCCATATGGCGGATCAGCTCACAGA |
| *CAM2Y-RP* | CGGGATCCTCACTTAGCCATCATAACCTTCA |
| *PEPR1-CY FP* | GGAATTCCATATGATGTGCCTACGTCGTCGCA |
| *PEPR1-CY RP* | CGGGATCCTTACCGAACTGAATCAGAGGA |
| *PEPR2-CY FP* | GGAATTCCATATGATGCGGTGCAAAAGAGGAAC |
| *PEPR2-CY RP* | CGGGATCCCTAGTGAACTGAACCCGAAGTGCT |
| *pGEMHE-CNGC13 FP1* | GGATCCATGGCTTTCGGAAG |
| *pGEMHE-CNGC13 RP1* | CACCAGCTCGCCTCTTCC |
| *pGEMHE-CNGC13 FP2* | GCTAGCGTGCTGAGGACATT |
| *pGEMHE-CNGC13 FP3* | GCGCGAAGCTTGCGAGAAG |
| *pGEMHE-CNGC13 FP4* | GAGGCACTTCTGTCTGGACC |
| *pGEMHE-CNGC13 FP5* | GGCGCACATGGGGAGCTA |
| *pGEMHE-CNGC13 FP6* | GTGCTTCGCTAGATACCAGA |
| *pGEMHE-CNGC13 FP7* | CATTTCTGGACTGGTGCTGTTT |
| *pGEMHE-CNGC13 RP2* | AAGGGCCCATGTCAGCAGAT |
| *CNGC13_nLuc_FP* | GGGGTACCATGGCTTTTGGCCGCAACAATC |
| *CNGC13_nLuc_RP* | ACGCGTCGACTTAAGGGTTTCGGAGACTG |
| *PEPR2_cLuc_FP* | CGGGATCCATGAGGAATCTTGGGTTACTCGA |
| *PEPR2_cLuc_RP* | ACGCGTCGACGTGAACTGAACCCGAAGTGCT |
| GCaMP3_FP | CCCCCCGGGATGGGTTCTCATCATCATCA |
| GCaMP3_RP | CCGCTCGAGTTACTTCGCTGTCATCATTTGT |

**Genotyping primers**

| *CNGC13 S_082668 LP* | GGTTGTTGTTTGCTTGCTTTC |
| --- | --- |
| *CNGC13 S_082668 RP* | AAACCCCACCAGAAGCAGTAG |
| *CNGC19 S _129200 LP* | AGGAGGGTGAGAGAGGTTGAG |
| *CNGC19 S _129200 RP* | CCAAGTTACTCAAGCCGAGTG |
| *CNGC19 S_27306 LP* | CGCGGATCTCTTTATTCACAC |
| *CNGC19 S_27306 RP* | ATGAGGATTCATTATTCCGGG |
| *CNGC13 S _057742 LP* | TTTAGGAGGTAGCAAAGGTAGGTTG |
| *CNGC13 S _057742 RP* | CAGCTATGCAATTCGAGAAGG |
| *LBb1.3* | ATTTTGCCGATTTCGGAAC |
| *COI1 FP* | GTTCTCTTTAGTCTTTAC |
| *COI1 RP* | CAGACAACTATTTCGTTACC |
| CNGC13gRNA FP1 | ATTGGCCGGTCCGGCCACCGTTTG |
| CNGC13gRNA RP1 | AAACCAAACGGTGGCCGGACCGGC |
| CNGC13gRNA FP2 | ATTGTTCAGAGATTGGATTTCAGA |
| CNGC13gRNA RP2 | AAACTCTGAAATCCAATCTCTGAA |
| *PKIR1.1 seq FP* | CACCGACTCGGTGCCAC |
| *PKIR1.1 seq RP* | CAGCTTACATTTTCTTGAAC |
