## Supplemental Figure for "The Ca^2+^ Channel CYCLIC NUCLEOTIDE GATED CHANNEL13 (CNGC13) regulates systemic wound signalling and immunity against Spodoptera herbivory"

### Slide 1
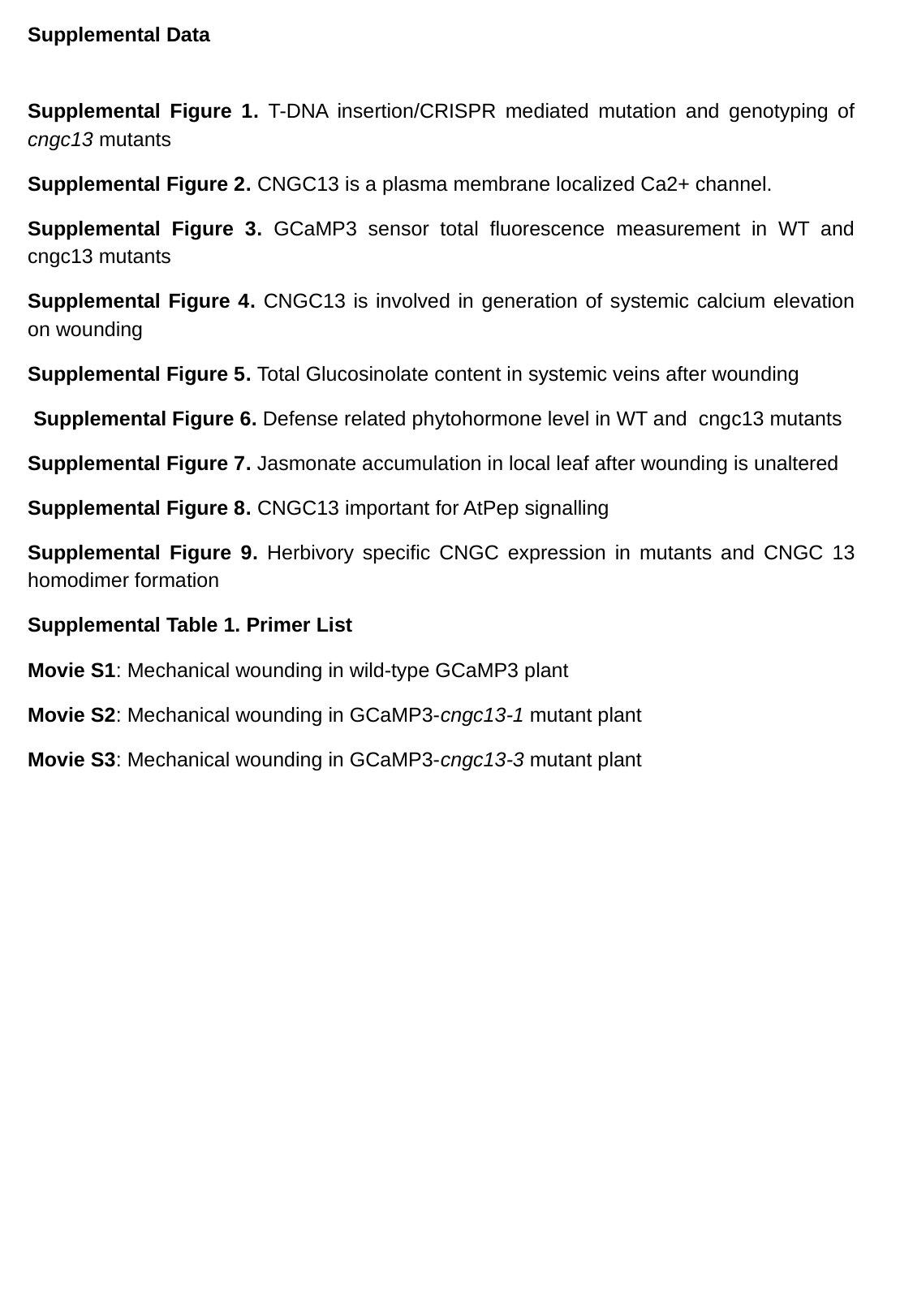

Supplemental Data
Supplemental Figure 1. T-DNA insertion/CRISPR mediated mutation and genotyping of cngc13 mutants
Supplemental Figure 2. CNGC13 is a plasma membrane localized Ca2+ channel.
Supplemental Figure 3. GCaMP3 sensor total fluorescence measurement in WT and cngc13 mutants
Supplemental Figure 4. CNGC13 is involved in generation of systemic calcium elevation on wounding
Supplemental Figure 5. Total Glucosinolate content in systemic veins after wounding
 Supplemental Figure 6. Defense related phytohormone level in WT and cngc13 mutants
Supplemental Figure 7. Jasmonate accumulation in local leaf after wounding is unaltered
Supplemental Figure 8. CNGC13 important for AtPep signalling
Supplemental Figure 9. Herbivory specific CNGC expression in mutants and CNGC 13 homodimer formation
Supplemental Table 1. Primer List
Movie S1: Mechanical wounding in wild-type GCaMP3 plant
Movie S2: Mechanical wounding in GCaMP3-cngc13-1 mutant plant
Movie S3: Mechanical wounding in GCaMP3-cngc13-3 mutant plant

### Slide 2
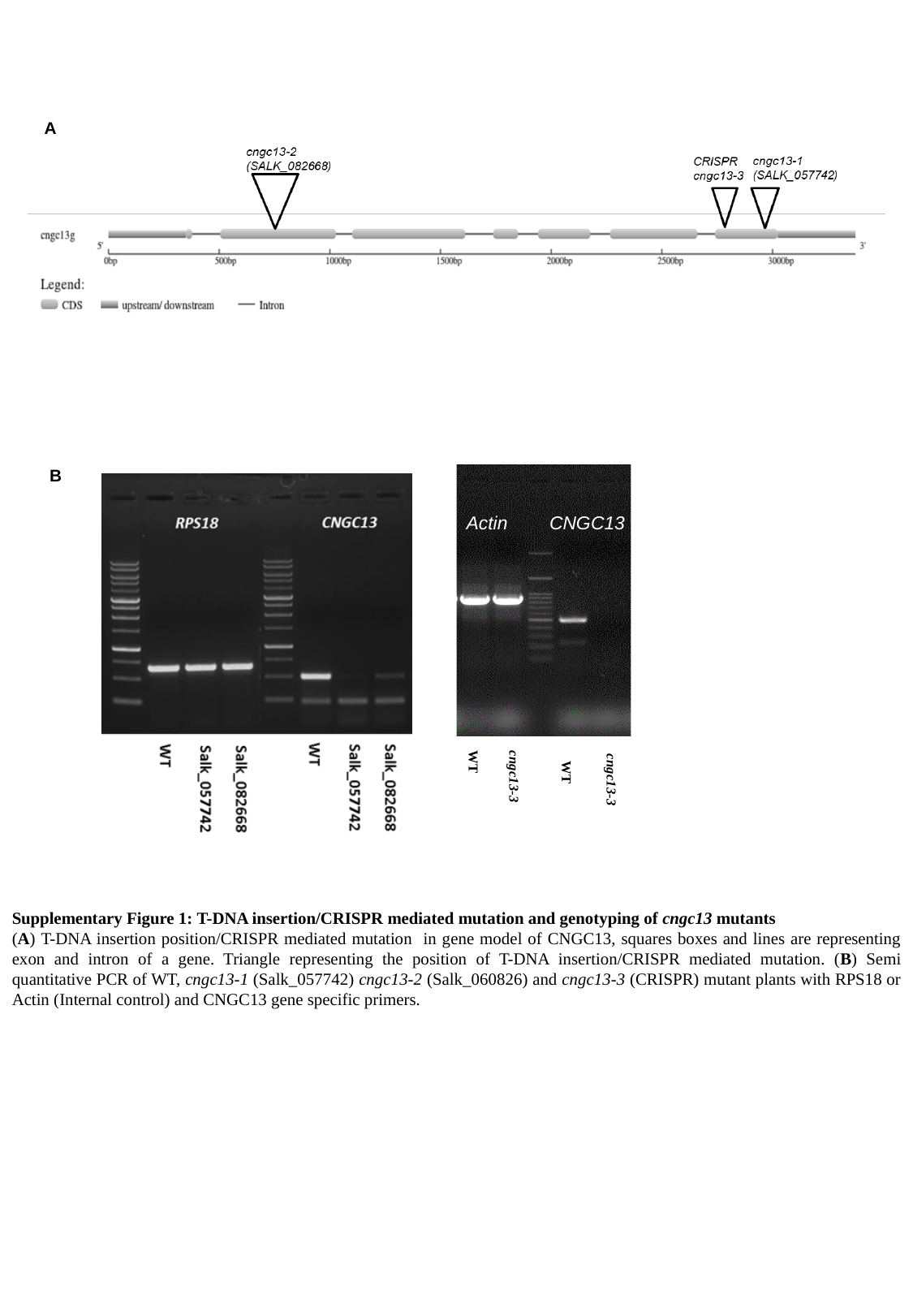

A
B
Actin
CNGC13
WT
WT
 cngc13-3
cngc13-3
Supplementary Figure 1: T-DNA insertion/CRISPR mediated mutation and genotyping of cngc13 mutants
(A) T-DNA insertion position/CRISPR mediated mutation in gene model of CNGC13, squares boxes and lines are representing exon and intron of a gene. Triangle representing the position of T-DNA insertion/CRISPR mediated mutation. (B) Semi quantitative PCR of WT, cngc13-1 (Salk_057742) cngc13-2 (Salk_060826) and cngc13-3 (CRISPR) mutant plants with RPS18 or Actin (Internal control) and CNGC13 gene specific primers.

### Slide 3
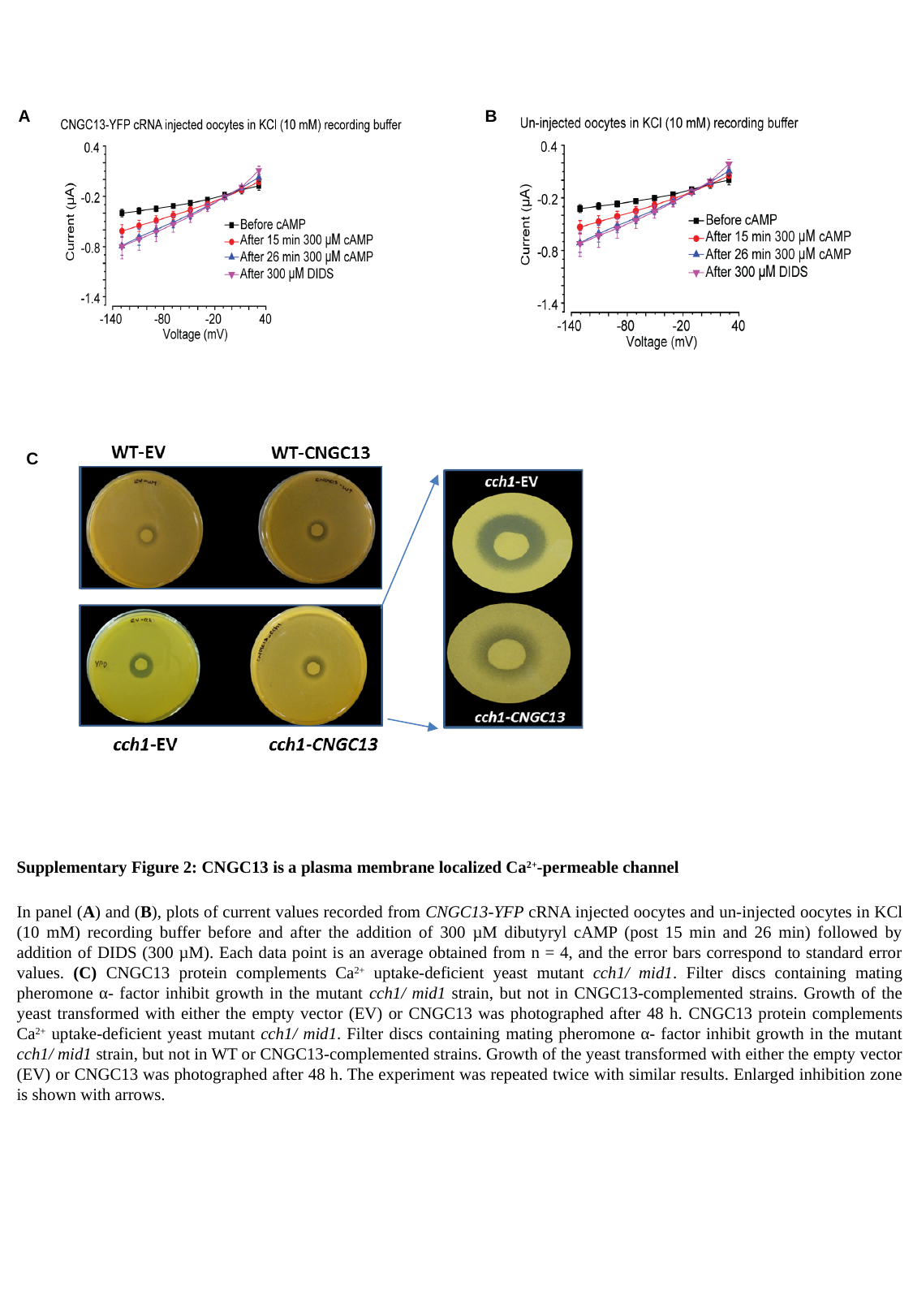

A
B
C
Supplementary Figure 2: CNGC13 is a plasma membrane localized Ca^2+^-permeable channel
In panel (A) and (B), plots of current values recorded from CNGC13-YFP cRNA injected oocytes and un-injected oocytes in KCl (10 mM) recording buffer before and after the addition of 300 µM dibutyryl cAMP (post 15 min and 26 min) followed by addition of DIDS (300 µM). Each data point is an average obtained from n = 4, and the error bars correspond to standard error values. (C) CNGC13 protein complements Ca^2+^ uptake-deficient yeast mutant cch1/ mid1. Filter discs containing mating pheromone α- factor inhibit growth in the mutant cch1/ mid1 strain, but not in CNGC13-complemented strains. Growth of the yeast transformed with either the empty vector (EV) or CNGC13 was photographed after 48 h. CNGC13 protein complements Ca^2+^ uptake-deficient yeast mutant cch1/ mid1. Filter discs containing mating pheromone α- factor inhibit growth in the mutant cch1/ mid1 strain, but not in WT or CNGC13-complemented strains. Growth of the yeast transformed with either the empty vector (EV) or CNGC13 was photographed after 48 h. The experiment was repeated twice with similar results. Enlarged inhibition zone is shown with arrows.

### Slide 4
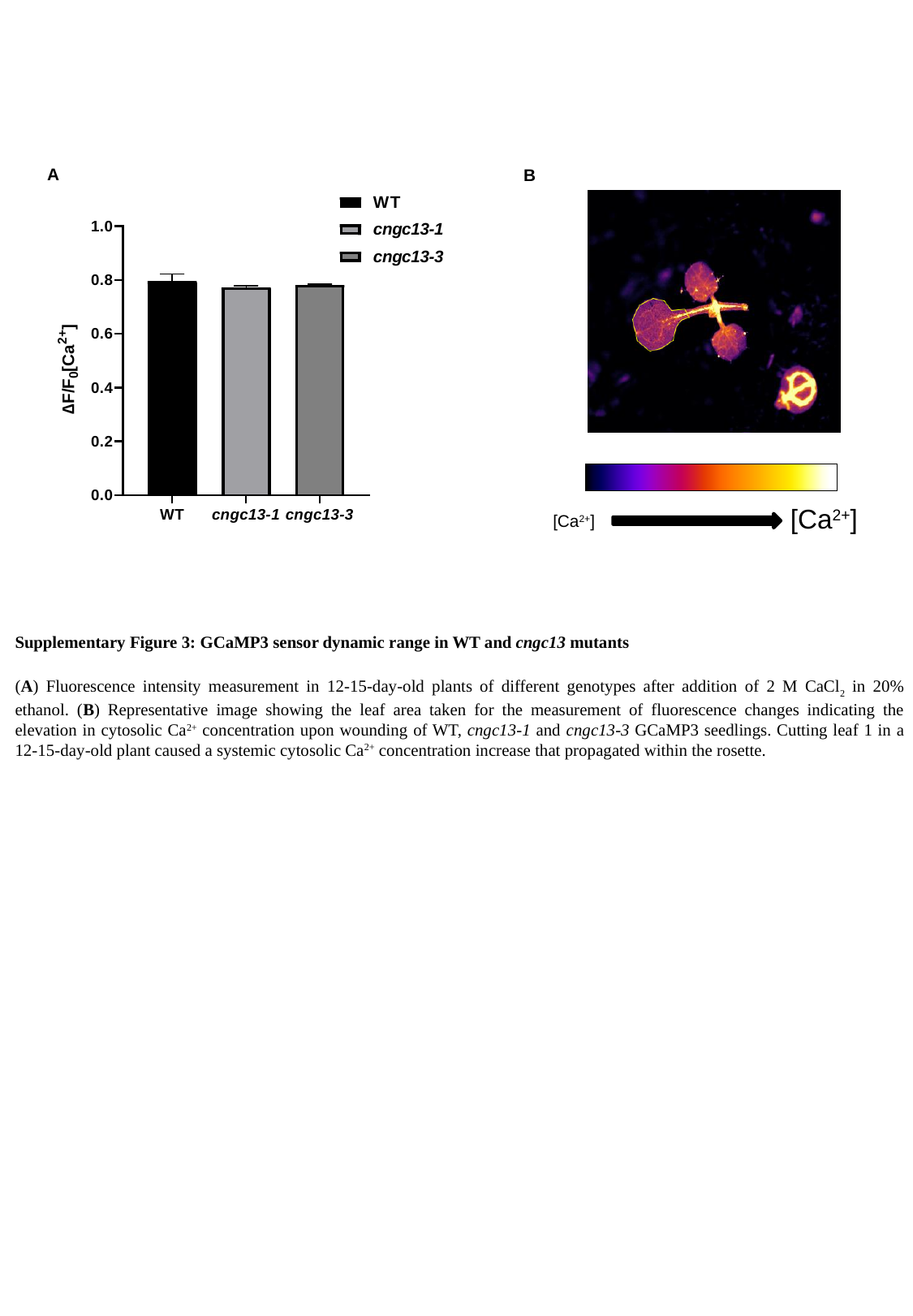

A
B
[Ca^2+^]
[Ca^2+^]
Supplementary Figure 3: GCaMP3 sensor dynamic range in WT and cngc13 mutants
(A) Fluorescence intensity measurement in 12-15-day-old plants of different genotypes after addition of 2 M CaCl_2_ in 20% ethanol. (B) Representative image showing the leaf area taken for the measurement of fluorescence changes indicating the elevation in cytosolic Ca^2+^ concentration upon wounding of WT, cngc13-1 and cngc13-3 GCaMP3 seedlings. Cutting leaf 1 in a 12-15-day-old plant caused a systemic cytosolic Ca^2+^ concentration increase that propagated within the rosette.

### Slide 5
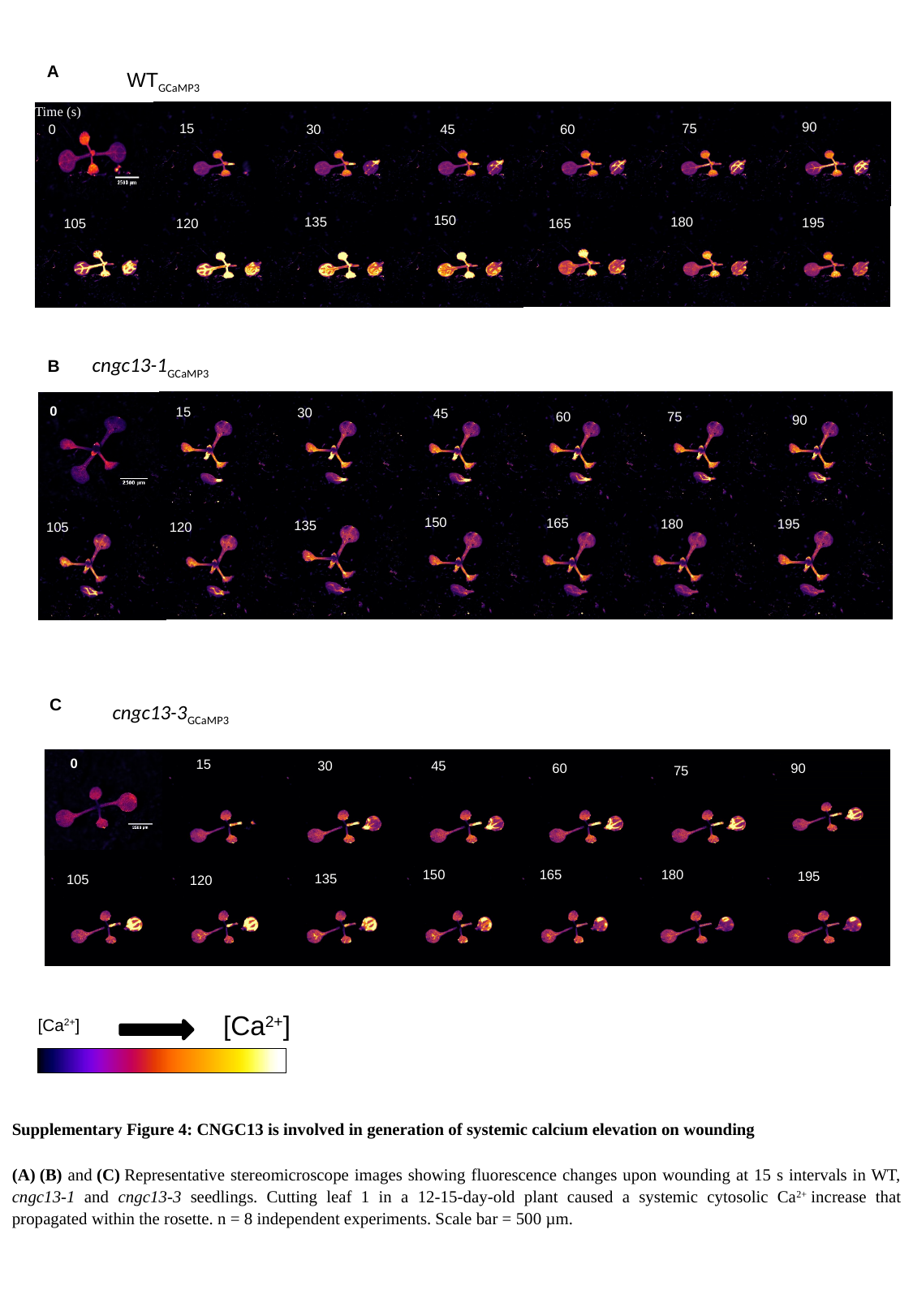

A
WT_GCaMP3_
Time (s)
90
15
75
30
45
60
0
150
135
180
195
105
120
165
cngc13-1_GCaMP3_
B
0
15
30
45
60
75
90
150
165
180
195
135
105
120
C
cngc13-3_GCaMP3_
0
15
30
45
60
90
75
150
165
180
195
135
105
120
[Ca^2+^]
[Ca^2+^]
Supplementary Figure 4: CNGC13 is involved in generation of systemic calcium elevation on wounding
(A) (B) and (C) Representative stereomicroscope images showing fluorescence changes upon wounding at 15 s intervals in WT, cngc13-1 and cngc13-3 seedlings. Cutting leaf 1 in a 12-15-day-old plant caused a systemic cytosolic Ca^2+^ increase that propagated within the rosette. n = 8 independent experiments. Scale bar = 500 µm.

### Slide 6
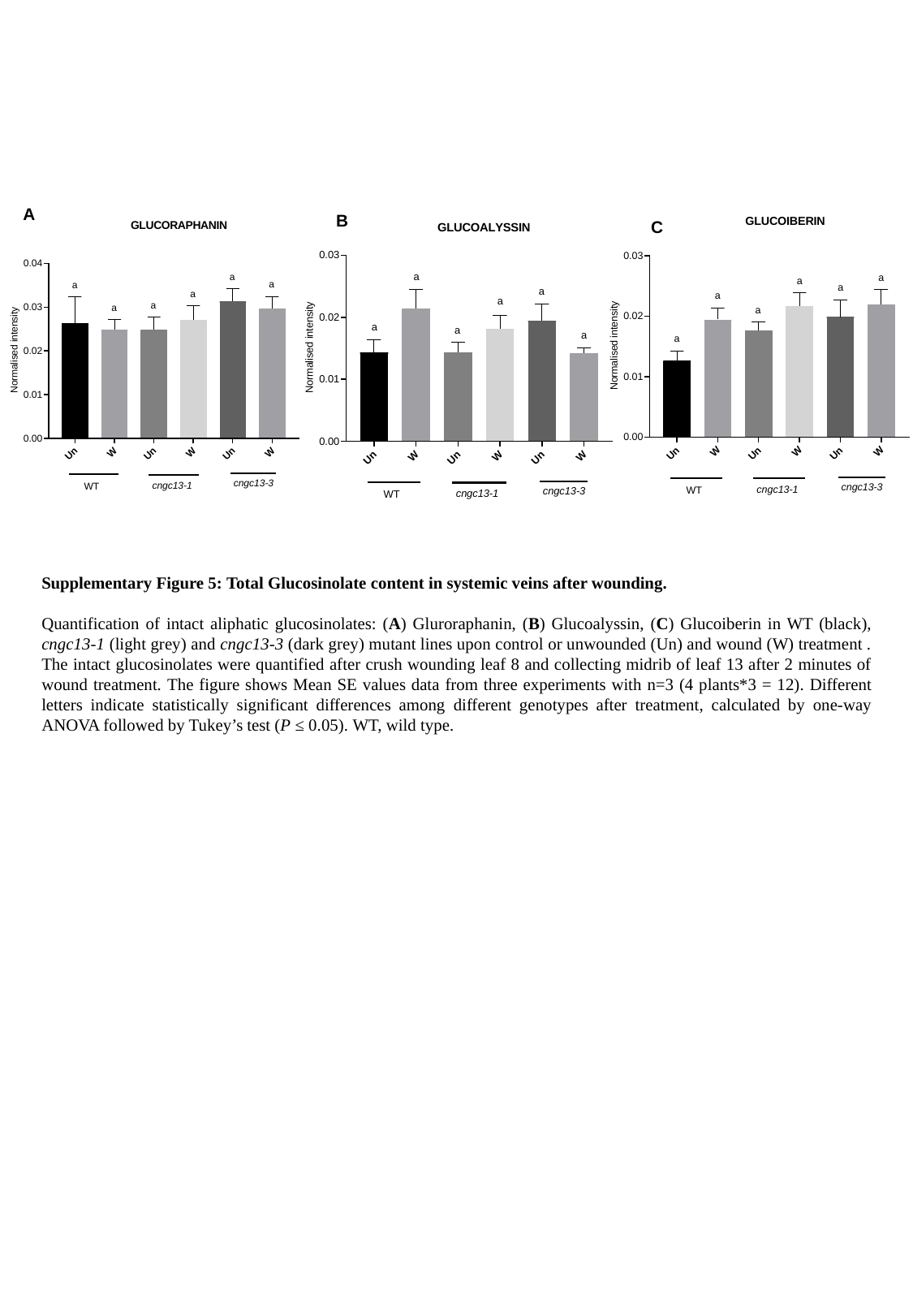

A
B
C

### Slide 7
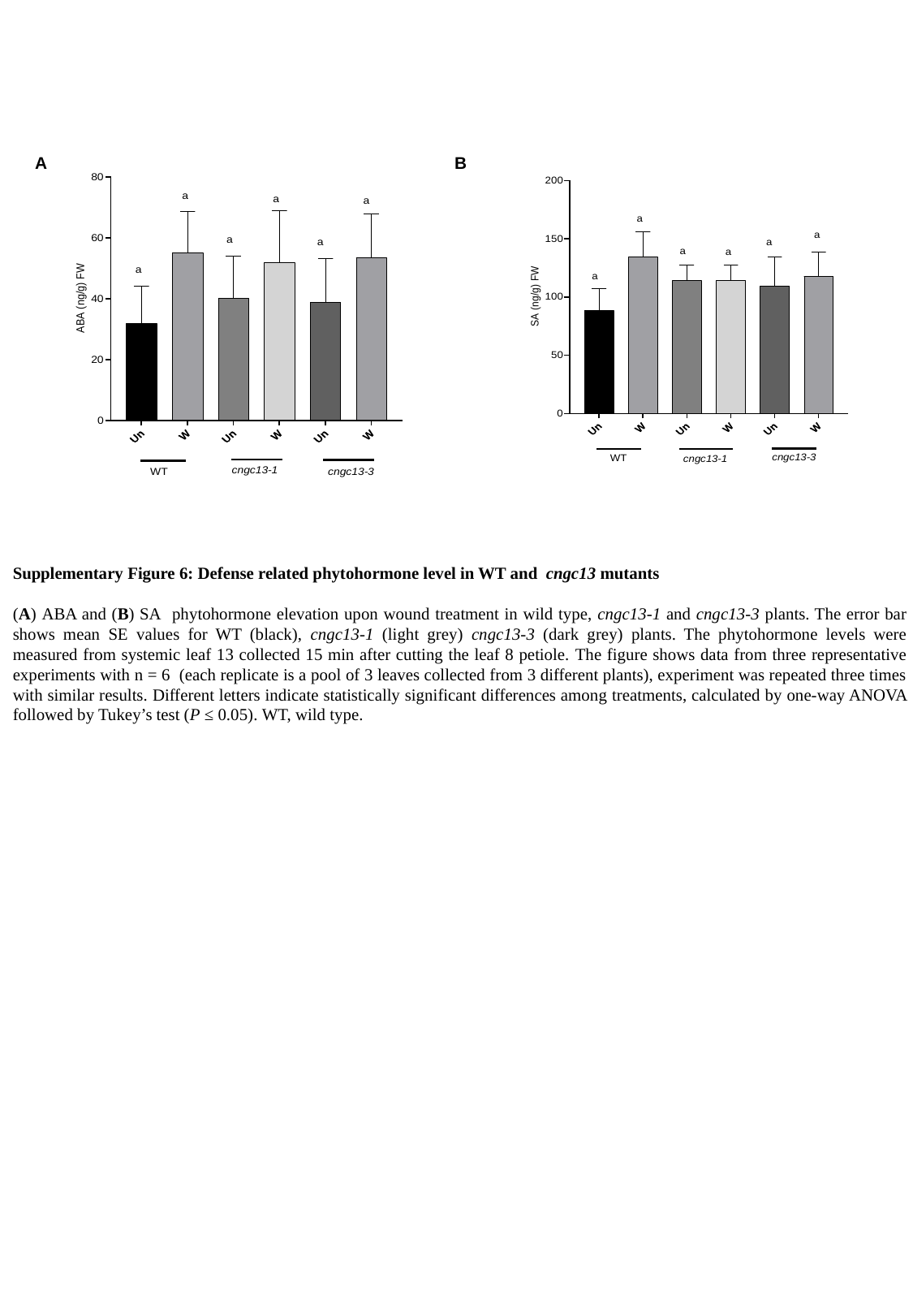

A
B

### Slide 8
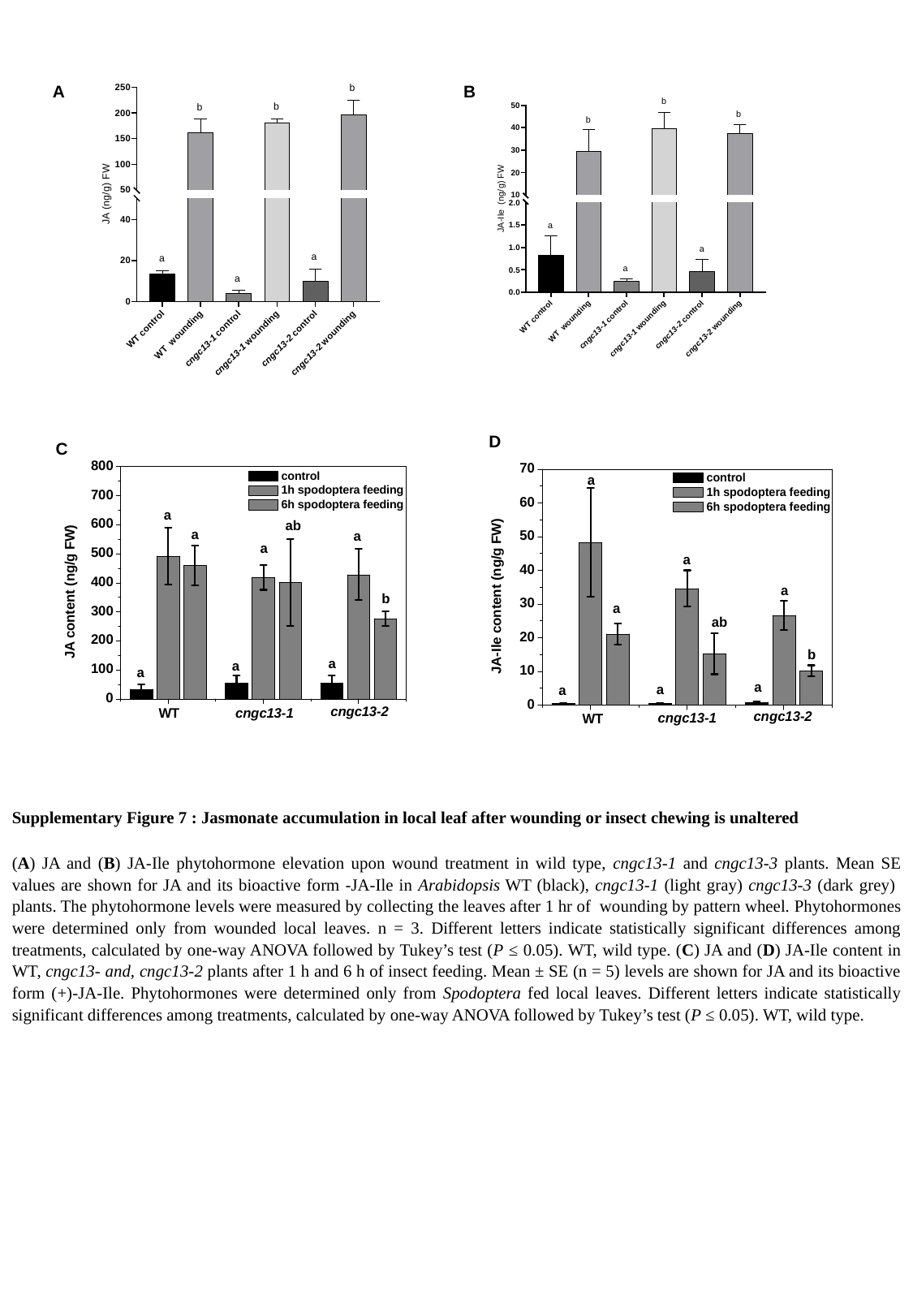

A
B
D
C

### Slide 9
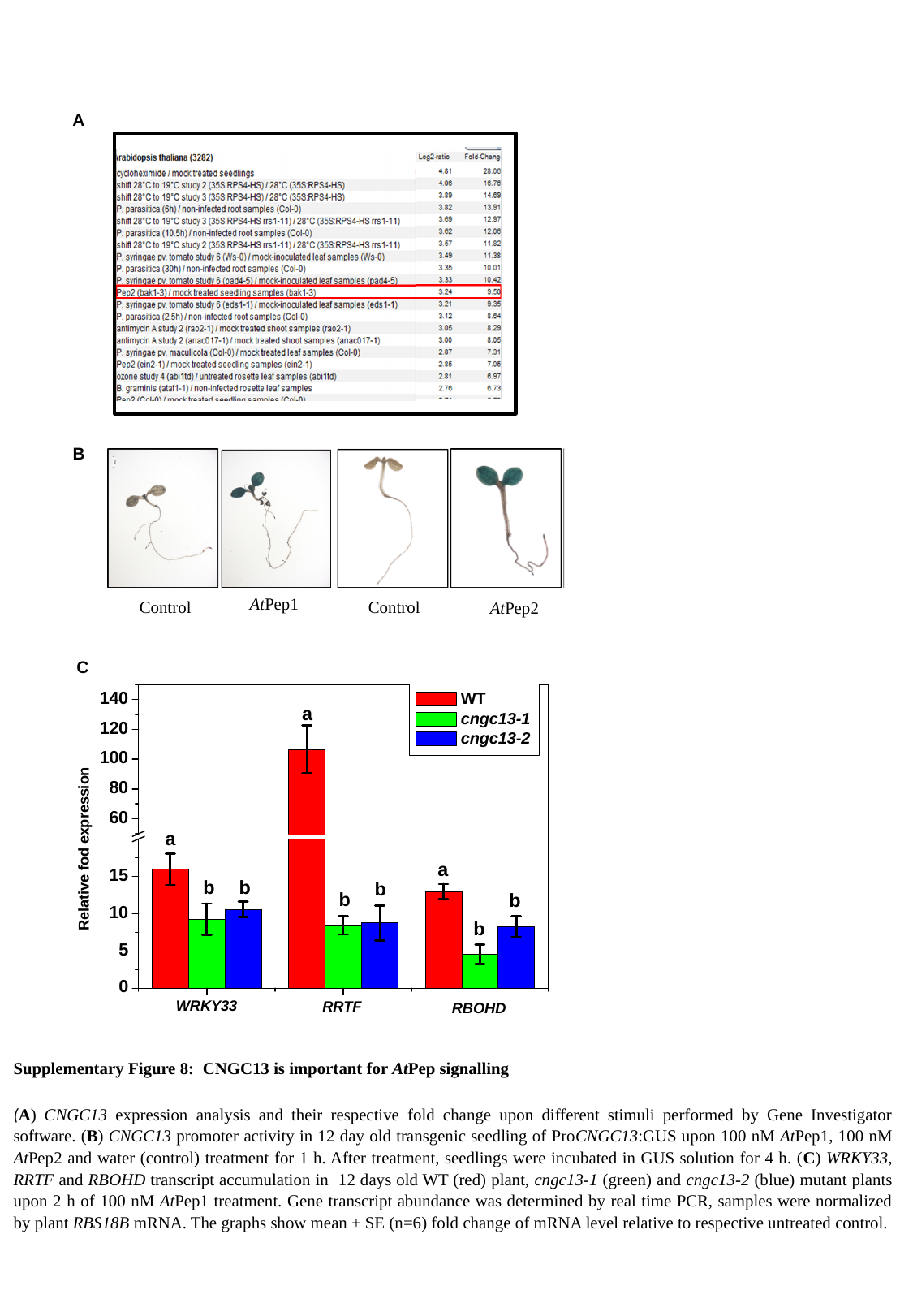

A
B
AtPep1
Control
Control
AtPep2
C
Supplementary Figure 8: CNGC13 is important for AtPep signalling
(A) CNGC13 expression analysis and their respective fold change upon different stimuli performed by Gene Investigator software. (B) CNGC13 promoter activity in 12 day old transgenic seedling of ProCNGC13:GUS upon 100 nM AtPep1, 100 nM AtPep2 and water (control) treatment for 1 h. After treatment, seedlings were incubated in GUS solution for 4 h. (C) WRKY33, RRTF and RBOHD transcript accumulation in 12 days old WT (red) plant, cngc13-1 (green) and cngc13-2 (blue) mutant plants upon 2 h of 100 nM AtPep1 treatment. Gene transcript abundance was determined by real time PCR, samples were normalized by plant RBS18B mRNA. The graphs show mean ± SE (n=6) fold change of mRNA level relative to respective untreated control.

### Slide 10
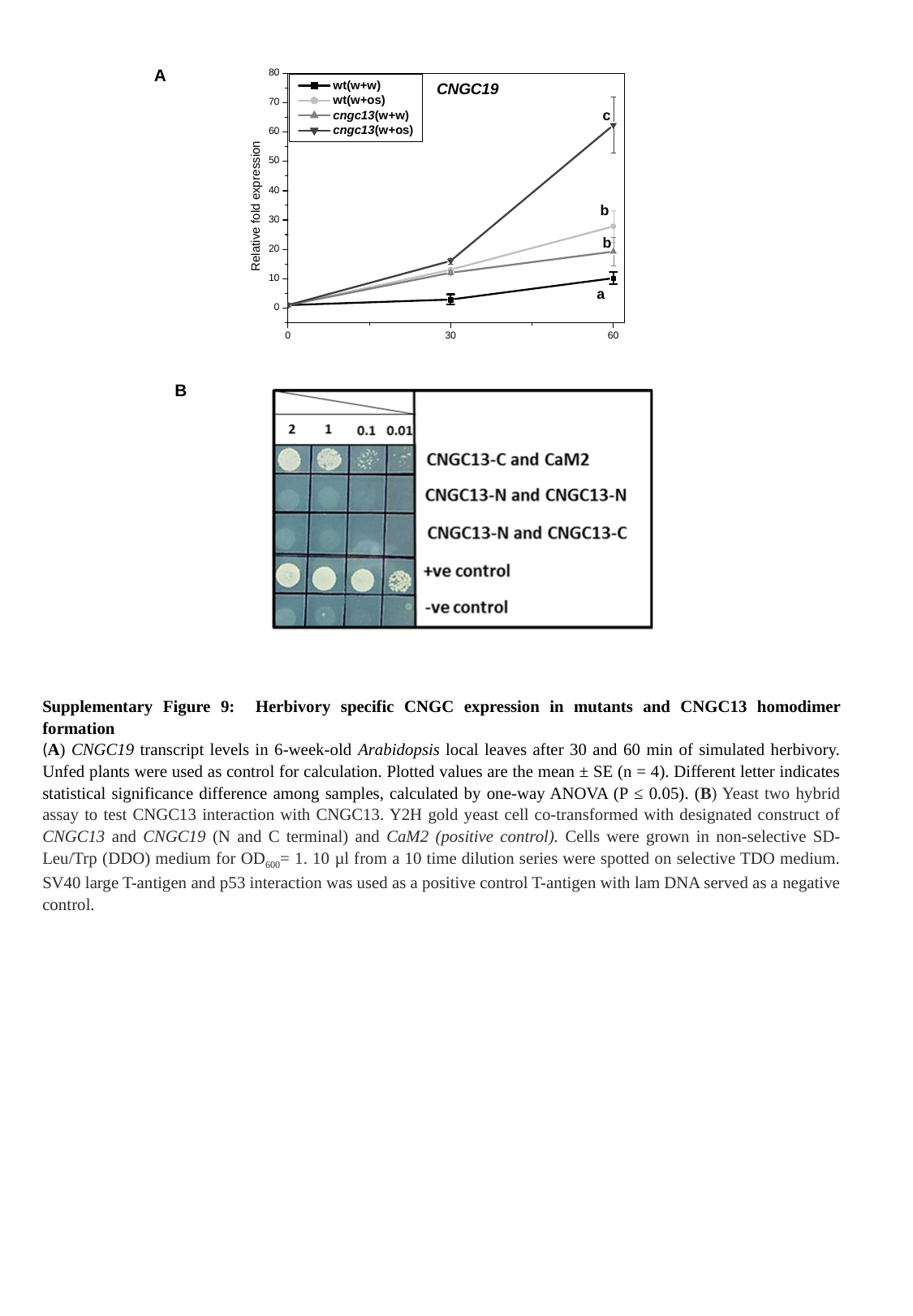

A
B
Supplementary Figure 9: Herbivory specific CNGC expression in mutants and CNGC13 homodimer formation
(A) CNGC19 transcript levels in 6-week-old Arabidopsis local leaves after 30 and 60 min of simulated herbivory. Unfed plants were used as control for calculation. Plotted values are the mean ± SE (n = 4). Different letter indicates statistical significance difference among samples, calculated by one-way ANOVA (P ≤ 0.05). (B) Yeast two hybrid assay to test CNGC13 interaction with CNGC13. Y2H gold yeast cell co-transformed with designated construct of CNGC13 and CNGC19 (N and C terminal) and CaM2 (positive control). Cells were grown in non-selective SD-Leu/Trp (DDO) medium for OD_600_= 1. 10 µl from a 10 time dilution series were spotted on selective TDO medium. SV40 large T-antigen and p53 interaction was used as a positive control T-antigen with lam DNA served as a negative control.

### Slide 11
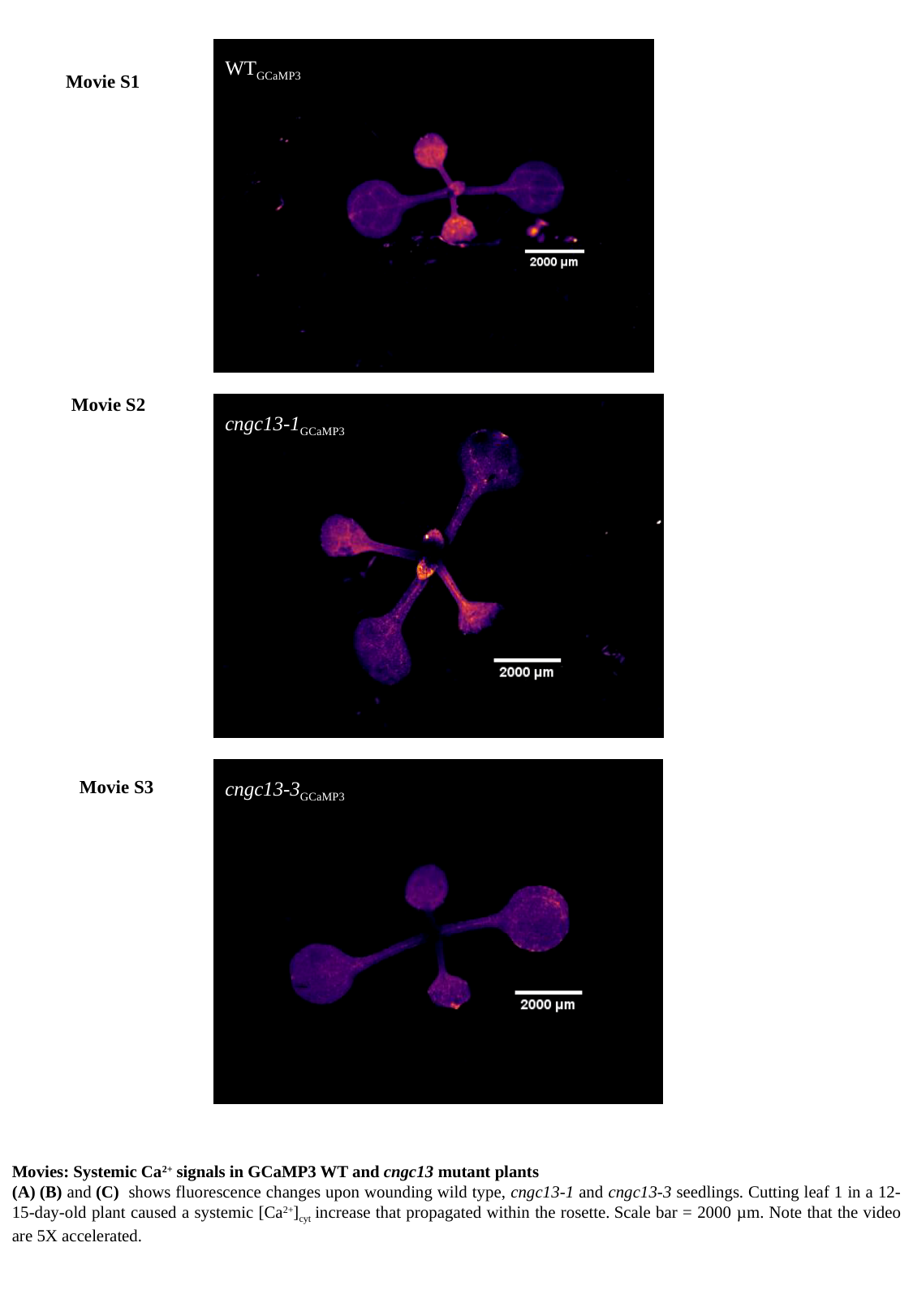

WT_GCaMP3_
Movie S1
Movie S2
cngc13-1_GCaMP3_
Movie S3
cngc13-3_GCaMP3_
Movies: Systemic Ca^2+^ signals in GCaMP3 WT and cngc13 mutant plants
(A) (B) and (C)  shows fluorescence changes upon wounding wild type, cngc13-1 and cngc13-3 seedlings. Cutting leaf 1 in a 12-15-day-old plant caused a systemic [Ca^2+^]_cyt_ increase that propagated within the rosette. Scale bar = 2000 µm. Note that the video are 5X accelerated.
